# Differential activation of JAK-STAT signaling in blood cell progenitors reveals functional compartmentalization of the *Drosophila* lymph gland

**DOI:** 10.1101/2020.07.26.219717

**Authors:** Diana Rodrigues, Yoan Renaud, K. VijayRaghavan, Lucas Waltzer, Maneesha S. Inamdar

**Author notes:** Author contributions: DR, KVR, LW and MSI conceptualized project and designed research; DR, LW and MSI performed experiments; DR, YR, LW and MSI analyzed data; DR, LW, KVR and MSI wrote the manuscript; LW and MSI obtained funding and facilities for the work.

## Abstract

Blood cells arise from diverse pools of stem and progenitor cells. Understanding progenitor heterogeneity is a major challenge. The *Drosophila* larval lymph gland is a well-studied model to understand blood progenitor maintenance and recapitulates several aspects of vertebrate hematopoiesis. However in-depth analysis has focused on progenitors located in lymph gland anterior lobes (AP), ignoring the progenitors from the posterior lobes (PP). Using *in situ* expression mapping and transcriptome analysis we reveal PP heterogeneity and identify molecular-genetic tools to study this abundant progenitor population. Functional analysis shows that PP resist differentiation upon immune challenge, in a JAK-STAT-dependent manner. Upon wasp parasitism, AP downregulate JAK-STAT signaling and form lamellocytes. In contrast, we show that PP activate STAT92E and remain undifferentiated. *Stat92E* knockdown in PP or genetically reducing JAK-STAT signaling permits PP lamellocyte differentiation. We discuss how heterogeneity and compartmentalization allow functional segregation in response to systemic cues and could be widely applicable.

**Highlights:** We provide an *in situ* and transcriptome map of larval blood progenitors Posterior lymph gland progenitors are refractory to immune challenge STAT activation after wasp parasitism maintains posterior progenitors

## Introduction

Blood and immune cells derive from hematopoietic stem cells (HSCs) that were classically thought to be a homogeneous population generating a defined hierarchy of progenitors. Recent studies reveal that mammalian HSCs and progenitor populations have dynamic cell surface marker phenotype and proliferative ability and varying *in vivo* differentiation potential in response to external cues (Ema et al., 2014, Crisan and Dzierzak, 2016, Haas et al., 2018). Studying aspects of intrapopulation heterogeneity and its implication in conditions of immune stress will improve our understanding of native and emergency hematopoiesis. Owing to the conserved signaling pathways and transcriptional factors that regulate hematopoiesis, *Drosophila melanogaster* has emerged as a powerful model to study blood cell development and the innate cellular immune response.

Like vertebrates, *Drosophila* hematopoiesis occurs in spatially and temporally distinct phases (Tepass et al., 1994, Evans et al., 2003, Holz et al., 2003, Krzemien et al., 2010). The embryonic wave of hematopoiesis primarily gives rise to macrophage-like phagocytic circulating and sessile hemocytes. A second wave of hematopoiesis takes place in a specialized larval hematopoietic organ, the lymph gland (LG), located dorsally along the anterior cardiac tube (Lanot et al., 2001, Mandal et al., 2004, Grigorian and Hartenstein, 2013, Rugendorff et al., 1994). In third instar larvae, the mature lymph gland is composed of a pair of anterior or primary lobes in segment T3/A1, followed by 2-3 pairs of lobes referred to as the secondary, tertiary and quaternary lobes - collectively called the posterior lobes (Banerjee et al., 2019). Based on morphology and molecular marker analysis, primary lobes are compartmentalized into distinct zones. The posterior signaling center (PSC), a small group of cells at the posterior tip of the primary lobes, specifically expresses Antennapedia (Antp) and acts as a signaling niche. The medullary zone (MZ), close to the cardiac tube, consists of multi-potent progenitors and is identified by expression of *Drosophila* E-cadherin (DE-cad), the complement-like protein Tep4 and reporters for the JAK-STAT pathway receptor Domeless (Dome). The peripheral cortical zone (CZ) consists of differentiated blood cells, *i*.*e*. mainly phagocytic plasmatocytes identified by Nimrod C1 (NimC1/P1) expression and a few crystal cells that express Lozenge (Lz) and the prophenoloxidases (ProPO). In addition, intermediate progenitors (IZ) reside in the region between the MZ and the CZ; they are identified by the expression of *dome* reporter and early differentiation markers like Hemolectin (Hml) or Peroxidasin (Pxn) but lack the expression of late markers like P1 for plasmatocytes and Lz for crystal cells (Banerjee et al., 2019, Jung et al., 2005).

The presence of blood cell progenitors in the anterior lobes prompted intense investigations to unravel how their fate is controlled. During normal development, anterior progenitor (AP) proliferation, quiescence and differentiation are finely orchestrated by the interplay of various pathways. Notably AP maintenance is controlled by activation of the Hedgehog and AGDF-A pathways in the MZ in response to signaling from the PSC and the CZ respectively (Mandal et al., 2007, Baldeosingh et al., 2018, Mondal et al., 2011). Moreover, within the MZ, Reactive Oxygen Species (ROS) levels, Wingless signaling and Collier expression regulate AP differentiation (Owusu-Ansah and Banerjee, 2009, Sinenko et al., 2009, Benmimoun et al., 2015). Besides, AP fate is controlled by systemic cues and external stresses such as immune challenge (Krzemien et al., 2010, Khadilkar et al., 2017b). In particular, deposition of egg from the parasitoid wasp *Leptopilina boulardi* (*L*. *boulardi*) in the hemocoel triggers AP differentiation into lamellocytes and premature histolysis of the primary lobes (Lanot et al., 2001, Crozatier et al., 2004, Louradour et al., 2017, Benmimoun et al., 2015, Bazzi et al., 2018, Letourneau et al., 2016, Small et al., 2014). ROS-mediated activation of EGFR and Toll/NF-kB signaling pathways in the PSC and the consecutive down-regulation of the JAK-STAT pathway in the AP are essential for this cellular immune response (Makki et al., 2010, Louradour et al., 2017).

In contrast with the anterior lobes, little is known about the posterior progenitors (PP) present in the posterior lobes (Banerjee et al., 2019). The general view is that these lobes essentially harbor progenitors as initially suggested by the higher expression of DE-cadherin and the lack of expression of mature blood cell markers (Jung et al., 2005). Yet only few studies on progenitor differentiation in the primary lobe report on phenotypes in the secondary lobes (Owusu-Ansah and Banerjee, 2009, Dragojlovic-Munther and Martinez-Agosto, 2013, Khadilkar et al., 2017b, Hao and Jin, 2017, Zhang and Cadigan, 2017, Kulkarni et al., 2011, Benmimoun et al., 2015). These studies revealed that posterior lobes could also differentiate in genetic contexts where there is extensive premature differentiation in primary lobes, however a thorough analysis of the posterior LG lobes is lacking. In addition, depletion of *asrij*, a*rf1 or garz* and overexpression of *arf1GAP*, all show more severe phenotypes of hyperproliferation and premature differentiation in the posterior lobes compared to the primary lobe (Kulkarni et al., 2011, Khadilkar et al., 2014). On the other hand, *Stat92E* mutation causes premature progenitor differentiation in the primary lobe but not in the posterior lobes (Krzemien et al., 2007), suggesting important differences in regulation and function of these lobes as well as inherent differences within the progenitor pool. Along that line, a recent single-cell RNA sequencing analysis of the LG has identified novel hemocyte sub-populations and suggests a higher degree of blood cell progenitor heterogeneity than previously thought (Cho et al., 2020). However the posterior lobes were excluded from this study and positional information is not retained in such single-cell sequencing analysis. A systematic mapping of lymph gland progenitors *in vivo* is not available and will provide valuable spatial information about this progenitor pool. The small size and contiguous nature of the *Drosophila* hematopoietic organ allows simultaneous comprehensive assessment of progenitors across lobes, which is essential for understanding the dynamics of progenitor heterogeneity and function in response to local and systemic cues. Such analysis is currently not feasible in vertebrate hematopoiesis. Here we combined *in situ* expression analysis of the entire LG with differential RNA sequencing analysis of the anterior and the posterior lobes to map expression of known blood cell markers and to identify new progenitor markers. Furthermore, by assessing the response to immune challenge, we reveal the functional heterogeneity of the LG progenitors and we propose that differential regulation of the JAK-STAT pathway underlies the maintenance of PP in response to wasp infestation.

## Materials and Methods

### Fly stocks and genetics

*Drosophila* stocks were maintained under standard rearing conditions at 25°C, unless specified otherwise. *Canton-S* was used as the wild type reference strain. *dome-Gal4,UAS-2xEGFP, Tep4-Gal4, Ser- Gal4,UAS2xEYFP* (provided by Prof. Utpal Banerjee, University of California Los Angeles). *pcol85-Gal4, UASmCD8-GFP* (provided by Prof. Michele Crozatier, University of Toulouse). *MSNF9-mcherry, hhF4f- GFP* (provided by Prof. Robert Schulz, University of Notre Dame), *domeMESO-GFP* (provided by Prof. Tina Mukherjee, NCBS), *upd3-Gal4,UAS-GFP* (provided by Dr. Sveta Chakrabarti, Indian Institute of Science), *UAS-Stat92E-RNAi* (VDRC #43866), *UAS-dlp-RNAi* (NCBS Fly facility) (VDRC #10299), *UAS-Stat92E* (#F000750) (Bischof et al., 2013) (Fly ORF, Zurich ORFeome project), *e33c-Gal4, arj*^*9*^*/arj*^*9*^, (Kulkarni et al., 2011). Fly-FUCCI (BL55121), *G-trace* (BL28281), *10xSTAT-GFP* (BL26197), *10xSTAT-GFP* (BL26198), *5-HT1B-GFP* (BL60223), *apt-GFP* (BL51550), *CG31522-GFP* (BL64441), *CG6024-GFP* (BL64464), *col (GMR13B08)-Gal4* (BL48546), *dlp (GMR53G07)-Gal4* (BL46041), *dlp (GMR53E05)-Gal4* (BL48196), *dlp (0421-G4)-Gal4* (BL63306), *E(spl)mß-HLH-GFP* (BL65294), *netB-GFP* (BL67644), *netBtm-V5* (BL66880), *sns-GFP* (BL59801), *Tsp42Ee-GFP* (BL51558), *UAS-Lifeact-RFP* (BL58362), *Ubx(M3)-Gal4* (provided by Prof. Ernesto Sanchez-Herrero, CBMSO, University of Madrid). To generate the *Pxn-GFP* reporter line, regulatory region (dm6 chr3L:2629330-2629639) was cloned into the pH-stinger vector (DGRC #1018) and the corresponding transgenic lines were generated by standard P-element-mediated transformation into *w*^*1118*^ flies.

### Whole lymph gland sample preparation

Briefly, third instar larvae or staged pupae were washed in 1X PBS, placed dorsal side facing up and pinned at the anterior and posterior ends. Larvae were slit laterally and the cuticle was carefully cut along the edges, loosened and lifted away gently to expose viscera which were then removed. The entire LG, attached to the brain lobes in the anterior and flanking the dorsal vessel which was thus exposed, was washed and fixed in this preparation. Fixed hemi-dissected larvae were transferred to a 96-well dish and processed for staining. For mounting, stained hemi-dissected preparations were transferred to a cover slip dish, the whole LG was carefully separated from rest of the larval cuticle with fine scissors.

### Sample preparation and RNA-sequencing

Lymph glands from *w*^*1118*^ female third instar larvae were dissected in ice cold PBS, the anterior and the posterior lobes of the lymph gland were separated, transferred to an Eppendorf containing 10µl of RNAlater (FisherScientific #10564445) and frozen on dry ice. Independent biological triplicates were prepared for each condition. RNA extraction was performed using Arcturus PicoPure RNA kit (ThermoFisher #KIT0204). RNA samples were run on the Agilent Bioanalyzer to verify sample quality. Samples were converted to cDNA using Nugen’s Ovation RNA-Seq System (Catalogue # 7102-A01). Libraries were generated using Kapa Biosystems library preparation kit (#KK8201) and multiplexed libraries were sequenced on a 1×75 High output flow cell on the NextSeq550 platform (Illumina). Reads were filtered and trimmed to remove adapter-derived or low-quality bases using Trimmomatic and checked again with FASTQC. Illumina reads were aligned to *Drosophila* reference genome (dm6Ensembl release 70) with Hisat2. Read counts were generated for each annotated gene using HTSeq-Count. RPKM (Reads Per Kilobase of exon per Megabase of library size) values were calculated using Cufflinks. Reads normalization, variance estimation and pair-wise differential expression analysis with multiple testing correction was conducted using the R Bioconductor DESeq2 package. Heatmaps and hierarchical clustering were generated with “pheatmap” R package. Gene ontology enrichment analyses were performed using Genomatix. The RNA-seq data were deposited on GEO under the accession number GSE152416.

### Larval oral infection assay

Early third instar larvae were starved in empty vials for 2–3h, then placed in fly food vials containing banana pulp alone (uninfected control) or mixed with concentrated bacterial pellets (*E*. *coli* or *P*.*entemophila*) followed by incubation at 25°C. 10-12h post infection, larvae were washed in 70% ethanol, rinsed in water, dissected and processed for immunostaining. *E*.*coli* (Khadilkar et al., 2017a); *P*. *entemophila* (provided by Dr. Sveta Chakrabarti, Indian Institute of Science).

### Wasp parasitism assay

4-5 female wasps (*Leptopilina boulardi*, provided by Prof. Tina Mukherjee, NCBS) were introduced into vials containing 40-50 late second instar (about 65h AEL) *Drosophila* larvae for 3-4 hours. After wasp removal larvae were allowed to develop further before dissection at 2, 3 or 4 days post-parasitism for dissection and immunostaining. Throughout the experiment larvae were maintained at 25°C. Lamellocytes were counted manually on the basis of DAPI positive nuclei that showed lamellocyte-like morphology scored by Phalloidin or that expressed *MSNF9-mcherry*.

### Primers and *in situ* hybridization probes

For *in situ* hybridization, DIG-UTP labeled anti-sense RNA probes were used (Avet-Rochex et al., 2010). Required fragments were PCR-amplified from wild-type genomic DNA and used as template for preparation of DIG-labeled probe using T7 RNA Polymerase (Promega, USA) according to the manufacturer’s instructions. Concentration of probe to be used for *in situ* hybridization assay was optimized with help of dot blot. Amplicon sizes and primers used for PCR amplification are: *latran*: (775 bp) forward primer- 5’ CCACCCAGGGCAGCATGCTC 3’; reverse primer 5’ taatacgactcactatagggCCTATTGCGCTCATGGACAC 3’; *Tep1*: (769 bp) forward primer 5’ CCTTAGCCCTCAATCCGGCC 3’; *Tep1* reverse primer 5’ taatacgactcactatagggAACCGTCGTTACGTTTGTAG 3’. *Tep4* (802pb) forward primer 5’ CAGGGCAGAAGTTCAGAGGC 3’, reverse primer 5’ taatacgactcactatagggGTCCGCCAGCACCGGAATGG 3’. *Dystroglycan* (Dg) probe was generated using GH09323 cDNA (from DGRC) and SP6 RNA polymerase (Promega) for anti-sense transcription.

### Immunostaining, *in situ* hybridization and microscopy

Wandering third instar larvae and staged pupae were used for dissection from timed embryos. Immunostainings were as described before (Kulkarni et al., 2011). Images were captured with Zeiss LSM 510 confocal or Zeiss LSM 880 confocal microscopes and analyzed using Zen black processing software and ImageJ. For *in situ* hybridization, lymph glands were fixed in 4% paraformaldehyde for 15 min, washed three times 15 min each in PBST and pre-incubated for 1h at 60°C in Hybridization Buffer (HB: 50% Formamide, 2X SSC, 1mg/ml Torula RNA, 0.05mg/ml, Heparin, 2% Roche blocking reagent, 0.1% CHAPS, 5mM EDTA, 0.1% Tween 20). Lymph gland preparations were then incubated overnight at 60°C with DIG-labeled RNA probe, followed by incubation for 1h in HB and for 30 min in 50% HB-50% PBST at 60°C and three washes 15 min each in PBST. 1% Goat serum was used as blocking agent for 30 min, followed by incubation with sheep anti-DIG antibody conjugated to alkaline phosphatase (Roche, Switzerland) (1:1000) for 2h. Larvae were extensively washed in PBST and the *in situ* hybridization signal was revealed with NBT/BCIP substrate (Promega, USA). Lymph glands were mounted in 70% glycerol. Images were captured using Olympus IX70 bright field microscope. To visualize the anterior and posterior lobes, 2 to 3 pictures (or Z-stacks) were captured along the A-P axis and stitched manually to reconstitute the whole lymph gland.

### Antibodies

Mouse anti-Antennapedia (1:20, DSHB #4C3), rat anti-DE-cadherin (1:10, DSHB #DCAD2), mouse anti-Ubx (1:20; DSHB #FP3.38), mouse anti-Dlp (1:50, DSHB #13G8), mouse anti-Myospheroid (1:50, DSHB #CF.6G11), mouse anti-P1 antibody (1:100, kind gift from Prof. Istvan Ando, Biological Research Center of the Hungarian Academy of Sciences), rabbit anti-GFP (1:1000, Clinisciences #TP401), chick or rabbit anti-GFP (1:500, Molecular Probes Inc.) rabbit anti-V5 (1:1000, ThermoFisher #PA1-993), mouse anti- ProPO antibody (1:5, Bioneeds), rabbit anti-Asrij (1:50, (Kulkarni et al., 2011)). Phalloidin was conjugated to Alexa-488 or Alexa-568 or Alexa-633 and secondary antibodies were Alexa-488, Alexa-568 or Alexa- 633 conjugated (Molecular Probes, Inc.).

### Image processing and analysis

For representative images, projections were made from complete Z-stacks and stitched manually to reconstitute the whole lymph gland. For Phalloidin, medial slices were stitched manually to avoid interference with the cardiac tube. Images were processed uniformly for brightness and contrast using Adobe Photoshop Elements 14. White dotted lines indicate lymph gland lobe boundaries, yellow asterisks indicate pericardial cells. Complete Z-stacks were considered for all analysis. GFP mean fluorescence intensity was measured using ImageJ software. The area to be measured for each lobe was marked with the help of the select tool, followed by intensity measurements. 3D images were reconstructed for lymph gland lobes using complete Z-stacks with the help of IMARIS software. DAPI 11 positive nuclei were counted by using the spots module in IMARIS. For membrane and cytoplasmic markers, surface module was used for 3D rendering. Distance between the 3D surface and spot was defined, with the help of modules- Find spots closer to surface and Find spots away from surface; number of nuclei that were positive or negative for a particular marker were determined. Quantifications were performed for the primary, secondary and tertiary lobes individually.

### Statistical analysis and quantification

In all assays, control and test genotypes were analyzed in parallel. Each experiment was repeated independently at least three times. Graphs and statistical analyses were done by Graphpad Prism 8.0. Statistically significant differences were indicated by * *p*<0.05, ** *p*<0.01 and *** *p*<0.001 and ns indicates non-significant. Error bars represent standard error of mean (SEM).

## Results

### Mapping gene expression in the larval lymph gland

In third instar larvae, the lymph gland posterior lobes are separated from the primary lobes and from each other by pericardial cells. To shed further light on these poorly characterized lobes, we used a dissection method that preserves the whole lymph gland (see M&M) and we assessed their contribution as well as their composition. By quantifying the number of cells (DAPI^+^) present in the different lobes, we found that LG posterior lobes contain about twice the number of cells as compared to primary lobes, indicating that they significantly contribute to the larval hematopoietic system (Figure 1A). To better define the identity of the blood cells in the posterior lobes, we then analyzed the expression of well-characterized PSC, MZ and CZ markers. Consistent with previous reports, we observed that the *hedgehog* reporter line *hhF4f-GFP* (Tokusumi et al., 2010), the Gal4 lines for *Serrate* (*Ser-Gal4, UAS2xEYFP*) (Lebestky et al., 2003) and *collier* (c*ol-Gal4,UASmCD8-GFP*) (Crozatier et al., 2004) as well as the transcription factor Antennapedia (Antp) (Mandal et al., 2007) were expressed in the PSC. Whereas *hhF4f-GFP, Ser-Gal4, UAS2xEYFP* and Antp did not show any expression in posterior lobes (Figure 1C, D), *col-Gal4-UASmCD8-GFP* is highly expressed in ±40% of the cells in the tertiary lobe (Figure 1B). Immunostaining also revealed high levels of Col in the PSC and tertiary lobes, while lower levels are detected in the MZ and the secondary lobes as reported earlier (Figure S1A) (Benmimoun et al., 2015). To assess the distribution of progenitors, we analyzed MZ markers expression using Gal4 enhancer trap lines in *Thioester-containing protein 4* (*Tep4-Gal4,UASLifeact-RFP*) (Avet-Rochex et al., 2010) and *domeless* (*dome-Gal4,UAS2xEGFP*) (Jung et al., 2005) and immunostaining against *Drosophila* E-cadherin (DE-cadherin) (Jung et al., 2005). Besides their expression in the MZ of the primary lobes, *Tep4-Gal4, dome-Gal4* and DE-cadherin are expressed in most cells of the secondary lobes and some cell clusters in the tertiary and quaternary lobes (Figure 1E, F, G). Similar patterns were observed using RNA *in situ* hybridization against *Tep4* or GFP immunostaining against a Shg-GFP (DE-cadherin) endogenous fusion or the *domeMESO-GFP* reporter (Figure S1B-D). Differentiated cells reside primarily in the CZ and can be visualized by the expression of *Peroxidasin* (*Pxn-GFP*), an early marker of differentiation, NimC1 (P1) a plasmatocyte marker, or Prophenoloxidase (ProPO) and Hindsight/Pebbled (Hnt/Peb), two crystal cell markers. The analysis of the expression of these markers showed that differentiated cells are rare in the posterior lobes (Figure 1E, F, G and Figure S1E). Finally, cell cycle analysis using the Fly FUCCI system (Zielke et al., 2014) showed that *Tep4*-expressing PP, which are a major fraction of the progenitor pool, are in the S or G2/ early M phase, like the AP (Figure 1H). Thus our analysis provides conclusive evidence that the posterior lobes are made up almost entirely of undifferentiated progenitors and actually represent the main reservoir of LG progenitors in the third instar larvae. Additionally, we demonstrate heterogeneity of gene expression across lymph gland progenitor populations (Figure 1I).

**Figure 1:**
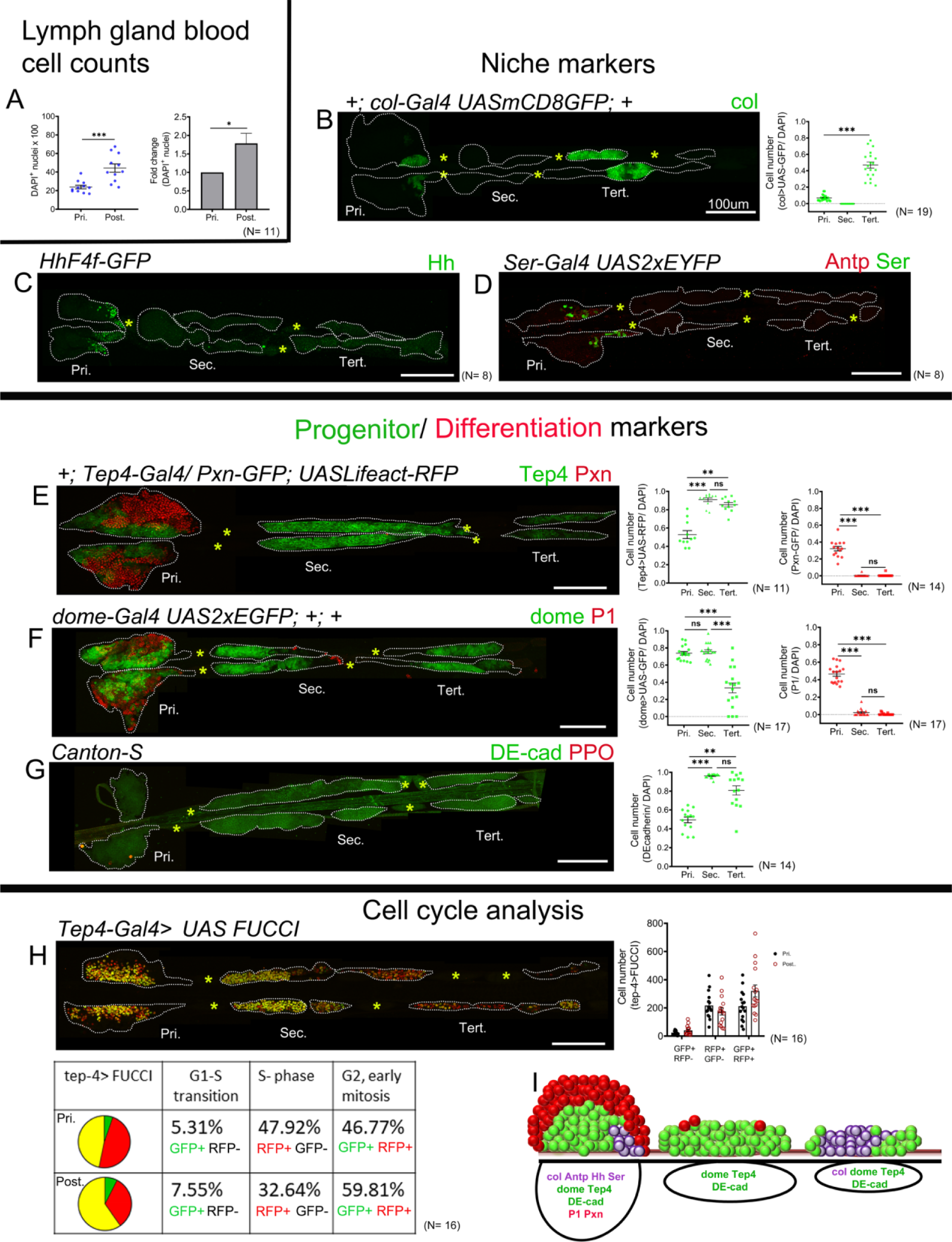
Gene expression analysis in the third instar larval lymph gland. Third instar larvae whole lymph gland preparations (including primary and posterior lobes). (A) Total LG blood cell counts (DAPI^+^ cells). (B-D) Expression profiles of PSC markers. (B) c*ol-Gal4,UASmCD8-GFP* (green) is expressed in the PSC and in the tertiary lobes; graph represents ratio of col>GFP^+^ /DAPI^+^ cells. (C) *hhF4f-GFP* (green) and (D) Antp (red) and *Ser-Gal4,UAS2xEYFP* (green) are expressed in the PSC only. (E-G) Expression profiles of MZ/progenitor (green) and CZ/differentiation (red) markers. (E) *Tep4-Gal4, UASLifeact-RFP* (green pseudo color), (F) *dome-Gal4,UAS2xEGFP* (green) and (G) DE-cadherin (green) expression is observed in the MZ as well as in the posterior lobes; (E) *Pxn-GFP* (red pseudo color), (F) P1 (red) and (G) ProPO (red) are restricted to the primary lobes. Quantification panels indicate ratios of Tep4^+^ /DAPI^+^ or Pxn^+^ /DAPI^+^ (E), dome^+^ /DAPI^+^ or P1^+^ /DAPI^+^ (F) and DE-cad^+^/ DAPI^+^ (G) cells respectively. (H) Cell cycle analysis for *Tep4-Gal4> FUCCI*. GFP+/RFP-: G1 phase; GFP-/RFP+: S-phase, GFP+/RFP+: G2/M phase. (I) Schematic representation of the expression of indicated markers in the third instar larval lymph gland. Pri. indicates primary lobes, Sec. indicates secondary lobes, Tert. indicates tertiary lobes and Post. indicates posterior lobes. (B-D, E-H) Yellow asterisks indicate pericardial cells. Lobes are outlined by white dashed lines. Nuclei were stained with DAPI, which is not displayed for clarity. (A) Mann-Whitney nonparametric test was used for analysis, (A) for fold change Student’s t-test was used for analysis. (B-G) Kruskal-Wallis test was used for statistical analysis. ** *p*<0.01, *** *p*<0.001, ns: non-significant and error bars represent SEM. (B-H) scale bar: 100µm.

### Anterior and posterior lobes of the larval lymph gland differ in their gene expression profile

To gain further insights into the antero-posterior compartmentalization of the larval lymph gland and to identify new blood cell markers, we established the transcriptome of the anterior and posterior lobes in wild type third instar wandering larvae (see materials and methods for details). Accordingly, the gene expression profile of manually-dissected anterior or posterior lobes was determined by RNA sequencing (RNA-seq) from biological triplicates using Illumina NextSeq550 sequencing system. We observed that 6709 genes (corresponding to ± 38% of the genes on *Drosophila* reference genome dm6) are expressed with a RPKM>1 in all 3 samples of the anterior lobes or of the posterior lobes (Table S1), including well known pan-hematopoietic markers such as *asrij* or *serpent* (*srp)*. Using DESeq2, we found that 406 genes are differentially expressed (p-value<0.01 & fold change >1.5) between the anterior and the posterior lobes, with 269 genes overexpressed in the anterior lobes and 137 genes overexpressed in the posterior lobes (Figure 2A and B, Table S2). In line with previous studies and the above results, markers of differentiated blood cells such as *Hemolectin* (*Hml), Nimc1, Pxn*, and *eater* for the plasmatocyte lineage, or *lz, PPO1, PPO2* and *peb*, for the crystal cell lineage, as well as PSC markers such as *Antp* and *Ser*, were overexpressed in the anterior lobes, whereas blood cell progenitor markers such as *DE-cadherin (shg), Tep4* or *col (col/kn)*, were overexpressed in the posterior lobes (Figure 2C). Gene ontology (GO) enrichment analyses showed a very strong over-representation for genes implicated in immune processes / defence responses and extracellular matrix (ECM) organisation among the genes overexpressed in the anterior lobes (Table 1 and Table S3); an observation consistent with the roles of differentiated blood cells in immune response and ECM synthesis (Banerjee et al., 2019). In contrast, GO analysis on the genes overexpressed in the posterior lobes were biased toward cell adhesion, neuronal differentiation and nephrocyte filtration (Table 1 and Table S3). While the latter GO enrichment is likely due to the presence of pericardial cells in dissected posterior lobes (see below), the neuronal link is unexpected and certainly warrants future investigations.

**Table 1.**
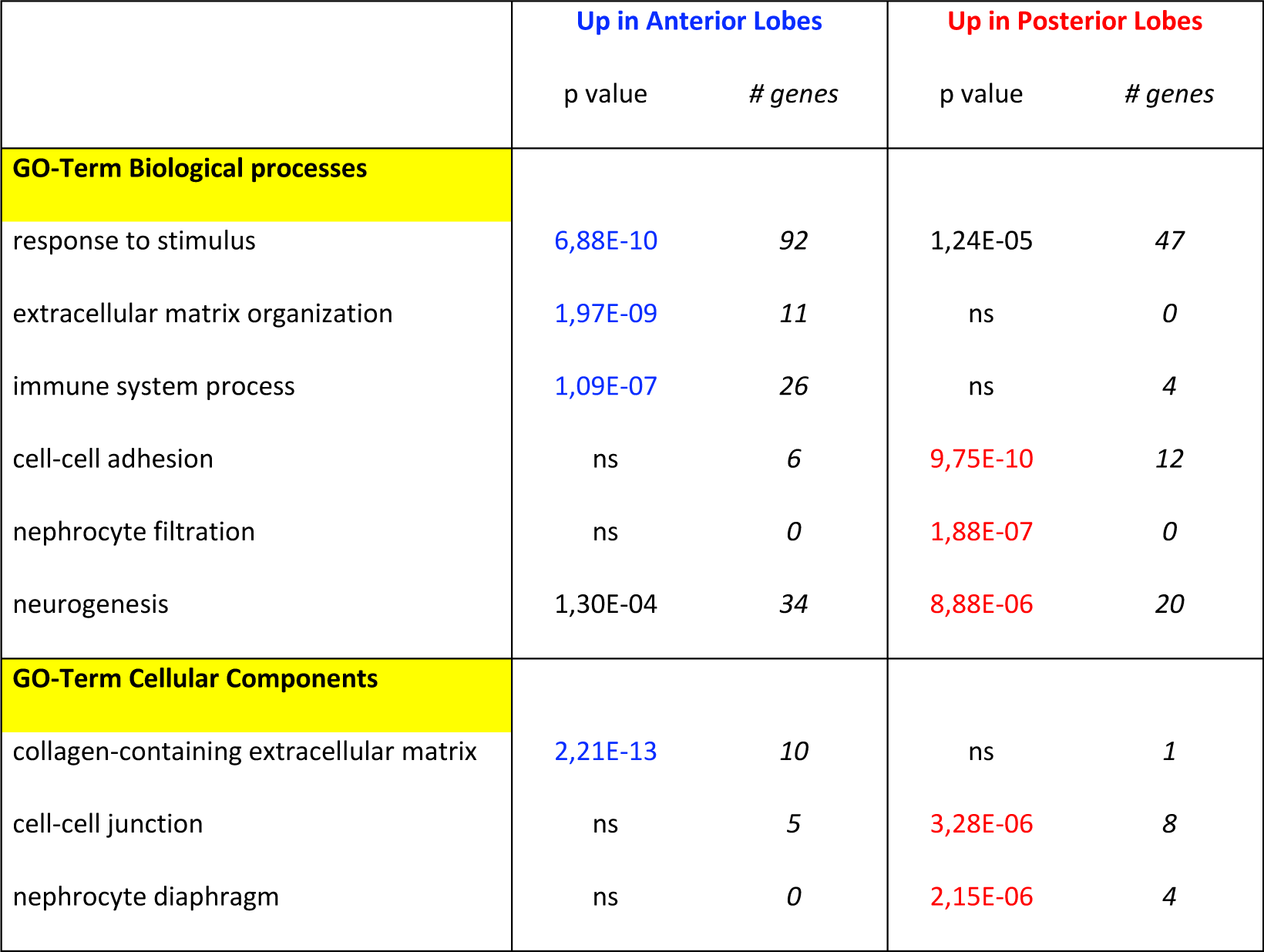
Main terms enriched in Gene Ontology analysis.

|  | Up in Anterior Lobes |  | Up in Posterior Lobes |  |
| --- | --- | --- | --- | --- |
|  | p value | # genes | p value | # genes |
| **GO-Term Biological processes** |  |  |  |  |
| response to stimulus | 6,88E-10 | 92 | 1,24E-05 | 47 |
| extracellular matrix organization | 1,97E-09 | 11 | ns | 0 |
| immune system process | 1,09E-07 | 26 | ns | 4 |
| cell-cell adhesion | ns | 6 | 9,75E-10 | 12 |
| nephrocyte filtration | ns | 0 | 1,88E-07 | 0 |
| neurogenesis | 1,30E-04 | 34 | 8,88E-06 | 20 |
| **GO-Term Cellular Components** |  |  |  |  |
| collagen-containing extracellular matrix | 2,21E-13 | 10 | ns | 1 |
| cell-cell junction | ns | 5 | 3,28E-06 | 8 |
| nephrocyte diaphragm | ns | 0 | 2,15E-06 | 4 |

**Figure 2:**
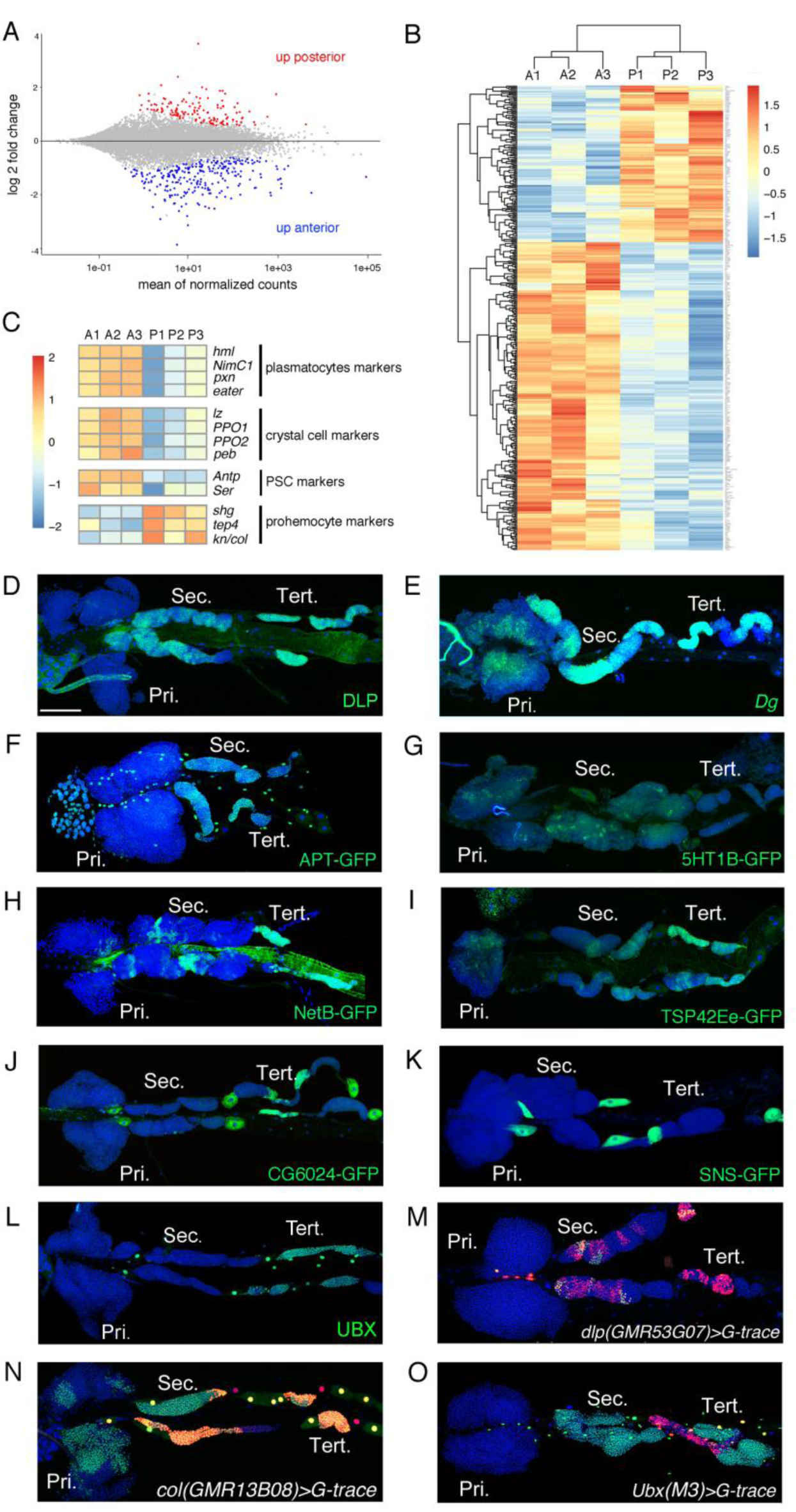
Characterization of the expression profile of the lymph gland anterior and posterior lobes and identification of new markers. (A) MA-plot of DESeq2 results for RNA-seq data comparison between anterior and posterior lymph gland lobes dissected from third instar larvae. Genes that are differentially expressed (adjusted p-value <0.01 and fold-change >1.5) are highlighted in red (up in posterior lobes) or blue (up in anterior lobes). (B) Heat map of differentially expressed genes between the anterior (A1, A2, A3) and posterior (P1, P2, P3) lobes RNA-seq samples. Hierarchical clustering was performed using R-Bioconductor. (C) Heat map of gene expression between anterior and posterior lobes for selected markers of lymph gland blood cell populations. (D-O) Third instar larvae whole lymph gland preparations showing the expression of the indicated genes or knock-in GFP fusions as revealed by immunostaining or RNA *in situ* (D-L) or the G-traced (green) and live (red) expression of the indicated Gal4 drivers (M-O). Nuclei were stained with DAPI (blue). Pri. indicates primary lobes, Sec. indicates secondary lobes, Tert. indicates tertiary lobes Scale bar: 100µm.

To validate our transcriptomic data and identify new markers for blood cell progenitors and/or the posterior lobes, we then analyzed the expression of several genes that were overexpressed in the posterior lobes according to our RNA-seq. Immunostaining against the heparan sulfate proteoglycan (HSPG) Dally-like protein (Dlp), which was described as a PSC marker (Pennetier et al., 2012) (see also Figure S2A), revealed that it is expressed at high levels in the posterior lobes and at very low levels in the MZ of the anterior lobes (Figure 2D). Consistent with the idea that its expression is activated by Col (Hao and Jin, 2017), Dlp displayed an antero-posterior gradient of expression similar to that of *col* (Figure 2D), with particularly strong levels in the tertiary lobes. RNA *in situ* hybridization against *Dystroglycan* (*Dg*), which encodes a cell surface receptor for the extracellular matrix, showed that it is strongly expressed throughout the posterior lobes and at lower level in the MZ of the anterior lobes (Figure 2E). Using an endogenously GFP-tagged version of the transcription factor Apontic (Apt), we found that it is expressed in the posterior lobes, in the heart tube, and at lower levels in the MZ (Figure 2F). Similarly, a GFP-5- hydroxytryptamine 1B (5-HT1B) fusion showed that this serotonin receptor is expressed in the posterior lobes and in the MZ (Figure 2G). GFP or V5 tagged versions of the guidance molecule Netrin-B (NetB) revealed that it is not only expressed throughout the cardiac tube but also in patches of cells in the posterior lobes, especially in the tertiary lobes, but not in the anterior lobes except for the PSC (Figure 2H and Figure S2B, D). Intriguingly, a GFP fusion with Tsp42Ee revealed that this Tetraspanin protein is strongly expressed in the posterior lobes, in particular in the tertiary lobes (Figure 2I). Furthermore, we found that the LDL receptor family member CG6024 was expressed in the tertiary lobes and in the PSC, as well as in the pericardial cells (Figure 2J and Figure S2C). Actually, 2 candidates (the Nephrin homolog Sticks and stone (Sns) and the fatty acid elongase CG31522) were specifically expressed in pericardial cells (Figure 2K and Figure S2E), indicating that contamination by these cells could contribute to the apparent overexpression of some genes in posterior lobe RNA-seq samples. Finally, immunostaining against Ultrabithorax (Ubx) showed that this HOX transcription factor is expressed in some pericardial and heart cells but only in the tertiary lobes within the lymph gland (Figure 2L). All together, these data indicate that the lymph gland prohemocytes are more heterogenous in terms of gene expression than previously thought.

In parallel, we used the G-trace system (Evans et al., 2009) to assess whether different Gal4 whose expression is under the control of putative enhancers of genes overexpressed in the posterior lobes can drive gene expression in specific territories of the lymph gland, and especially in progenitors and/or in the posterior lobes. Accordingly, we identified three Gal4 lines placed under the control of *dlp* regulatory regions that drive expression in part of the tertiary and secondary lobes (Figure 2M and Figure S2F, G). We also identified an enhancer in *col* driving expression in the MZ as well as in most cells of the posterior lobes (Figure 2N), contrary to the classically used *pcol85* lines, which is restricted to the

PSC and part of the tertiary lobes (Benmimoun et al., 2015). Importantly we found that the *Ubx(M3)- Gal4*, which essentially reproduces *Ubx* expression (de Navas et al., 2006), is expressed throughout the posterior lobes but not in the anterior lobes (Figure 2O), indicating that the anterior and posterior lobes emerge from distinct territories. To the best of our knowledge, this is the first posterior lobe-specific driver identified and it could be very useful to manipulate gene expression in these cells.

### Heterogeneity in gene expression is maintained in the pupal lymph gland

Previous studies indicated that lymph gland lobes histolyze during the course of pupal development (Lanot et al., 2001, Grigorian et al., 2011). Secondary lobes histolyze by approximately 8h (hours) APF (after pupa formation) (Grigorian et al., 2011). However the analysis did not cover the entire LG or analyze progenitor marker expression. For a more comprehensive analysis of the fate of posterior lobes, we analyzed the whole LG in hemi-dissected pupal preparations at different time-points. Expectedly, at 0h APF LG are intact and by approximately 5h APF primary lobes histolyze. Secondary and tertiary lobes begin to histolyze by approximately 10h APF and by 12-15h APF most of the LG is histolyzed (Figure 3A-D). Interestingly *col* (c*ol-Gal4,UASmCD8-GFP)* expression is maintained in the tertiary lobes and both *dome* (*dome-Gal4,UAS2xEGFP*) and *Tep4* (*Tep4-Gal4,UASLifeact-RFP*) are expressed in most cells of the secondary and tertiary lobes at 5 and 10h APF (Figure 3E-J), indicating that progenitors could be maintained during pupal development. Consistent with this hypothesis, we found few differentiated cells at these time points in the posterior lobes as marked by P1 or *Pxn-GFP* (Figure 3G-J). Thus our analyses show that posterior progenitors are maintained up to at least 10h of pupal development.

**Figure 3:**
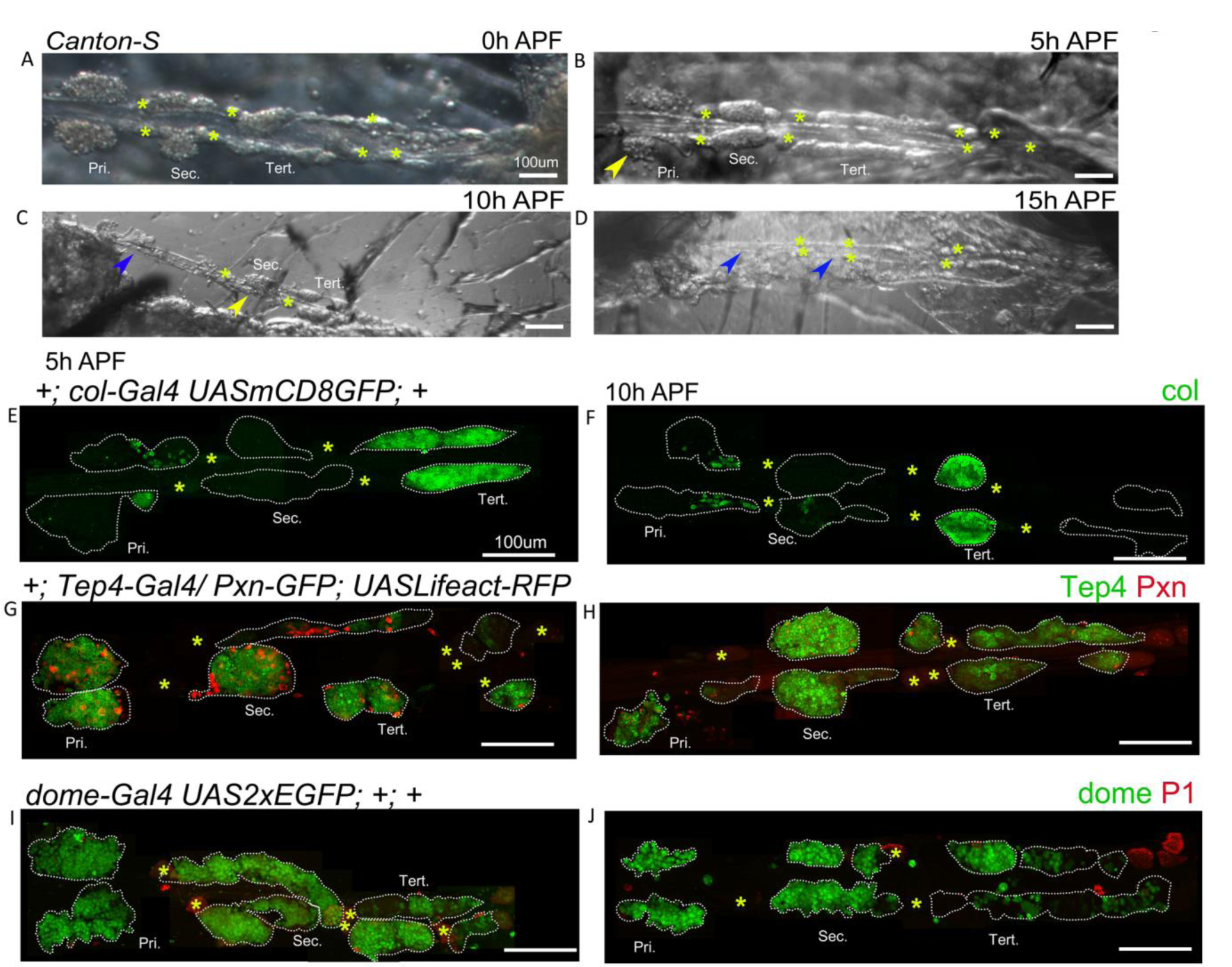
Gene expression analysis in the pupal lymph gland. (A-D) whole lymph gland hemi-dissected preparations indicating lobe histolysis during pupal development at 0h APF, 5h APF, 10h APF and 15h APF. (A-D) Yellow arrowheads indicate histolyzing lobes and blue arrowheads indicate histolyzed lobes. (E, F) c*ol-Gal4,UASmCD8-GFP* (green) is expressed in the PSC and tertiary lobes during 5h APF, 10h APF. (G, H) 5h APF, 10h APF lymph glands express *Tep4-Gal4> UASlifeact-RFP* (green pseudo color) in the primary lobes and posterior lobes; very few cells express *Pxn-GFP* (red pseudo color) in the primary and posterior lobes. (I, J) 5h APF, 10h APF lymph glands express *dome-Gal4,UAS2xEGFP* (green) in the primary lobes and posterior lobes, very few cells express P1 (red). (A-J) Yellow asterisks indicate pericardial cells, scale bars: 100µm. (E-J) nuclei were stained with DAPI (not shown for clarity). Pri. indicates primary lobes, Sec. indicates secondary lobes, Tert. indicates tertiary lobes.

### An anterior-posterior graded response to immune stress

*Drosophila* blood cells respond to immune stress caused by bacterial infections and wasp parasitism (Lanot et al., 2001, Sorrentino et al., 2002, Crozatier et al., 2004, Khadilkar et al., 2017b, Louradour et al., 2017, Sinenko et al., 2011). While all progenitors are assumed to respond uniformly to a given stress, especially to systemic cues, our findings that posterior progenitors differ in gene expression led us to hypothesize that this may reflect in the ability to maintain progenitors or differentiate upon immune challenge. Interestingly, infection with the Gram-negative bacteria *E*. *coli* promoted differentiation in the primary lobes as reported earlier (Khadilkar et al., 2017b), but not in the posterior lobes (Figure S3A-B).

*E*. *coli* has been used extensively to study the innate cellular immune response, however *E*. *coli* does not infect *Drosophila* naturally. *Drosophila* are natural host to parasitoid wasp and the hemocyte immune response is well-studied in this context (Banerjee et al., 2019). Hence we tested the sensitivity of the progenitor pools to parasitism by the specialist parasitoid wasp *Leptopilina boulardi*. Egg deposition by *L*. *boulardi* activates the humoral and cellular arms of immunity, leading to the production of lamellocytes, notably by the lymph gland, that encapsulate the wasp egg. The response to wasp infestation is well-characterized for the primary lobes but data for the posterior lobes is limited (Lanot et al., 2001). Of note too, the timing of response seems variable from lab to lab. For instance, Lanot *et al*. (2001) observed lamellocyte formation from 10h post-parasitism in the primary lobes, while Sorrentino *et al*. (2002) reported that lamellocyte production in the primary lobes begins around 51h post-parasitism and intensifies by 75h. Thus to assess lamellocyte differentiation and score for morphological changes in the posterior lobe progenitors, if any, we analyzed lymph glands of larvae at 3 days (approximately 75h) post-parasitism.

Phalloidin staining or Misshapen (using the *MSNF9-mCherry* reporter; (Tokusumi et al., 2009)) expression were used to detect lamellocytes. To account for the inter-individual variation in the extent of the response, we quantified the phenotypes based on the following broad classification – (a) Strong: lobes completely histolyzed and hence lamellocytes absent or few remnant cells attached to the cardiac tube, (b) Medium: few lamellocytes with histolyzing lobes that show uneven or discontinuous boundaries with loose packing of cells, (c) Mild: no lamellocytes but some cells in the lobe fuse or coalesce in groups with disrupted cell-cell boundaries (d) unaffected: lobes with undisturbed morphology, intact cell-cell connections and no lamellocytes.

Analysis of the LG post-parasitism revealed differences in the response of progenitors from anterior to posterior. In *Canton-S* (wildtype) background, 93.9% (46/49) of infested larvae showed some response in the primary lobes: 6.1% (3/49) had strong phenotypes, 85.7% (42/49) medium, and 2% (1/49) mild. Only 6.1% (3/49) of anterior lobes seemed unaffected. In contrast, 36.7% (18/49) of the larvae showed no effect on posterior lobes (Figure 4A, B). In addition, posterior lobes showed only mild phenotypes such as absence of clear cell-cell boundaries with occasional lamellocytes, but there was no strong phenotype and they remained intact. However, as reported before (Lanot et al., 2001, Sorrentino et al., 2002, Crozatier et al., 2004), posterior lobe size increased upon infestation, indicating that these lobes were indeed responding to the infection.

**Figure 4:**
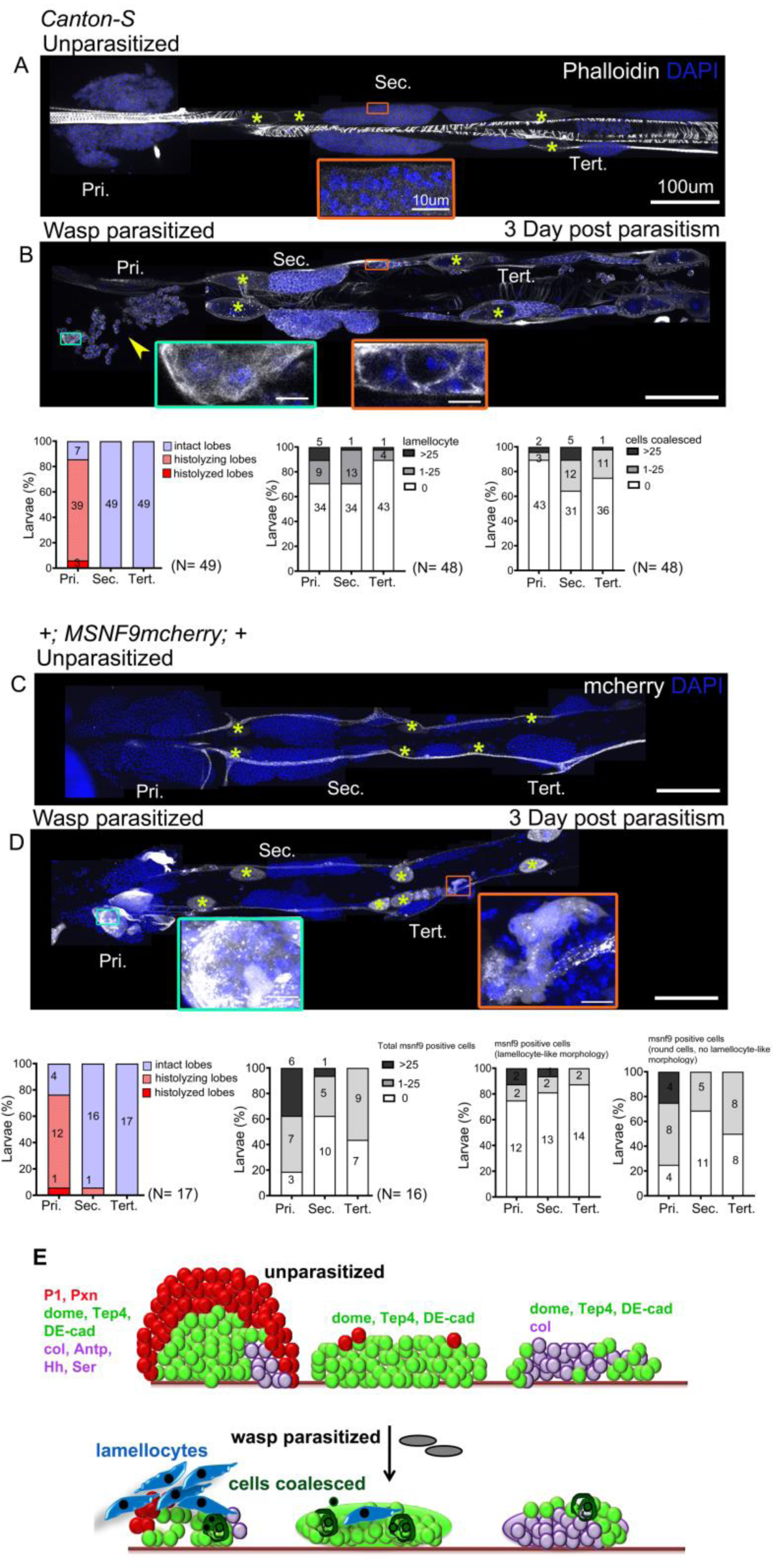
Lymph gland progenitor response to wasp parasitism. (A-B) *Canton-S* wild type strain lymph gland lobes were analyzed 3 days post-parasitism by *L*. *boulardi*. Age-matched unparasitized larvae were used as controls. Phalloidin (white) marks actin and was used for identifying lamellocytes as well as changes in cell morphology. (B) Yellow arrowhead indicate disintegrating primary lobes, green inset shows lamellocyte formation in the primary lobes and orange inset displays compromised cell boundaries in the secondary lobes. (C, D) *MSNF9-mcherry* larvae were analyzed for lamellocyte formation 3 days post-parasitism. Green and orange insets show lamellocytes in the primary and the posterior lobes respectively. For quantifications, nuclei showing lamellocyte morphology (phalloidin) or lamellocyte marker expression (MSNF9-mCherry, white pseudo color) were manually counted. Graphs indicate quantification for the cellular phenotypes: premature histolysis of lymph gland lobes, lamellocyte induction or compromised cell boundaries. Pri. indicates primary lobes, Sec. indicates secondary lobes, Tert. indicates tertiary lobes. Yellow asterisks indicate pericardial cells. (E) Schematic representation of the lymph gland response to wasp parasitism. Scale bars: 100µm or 10µm in inset.

To ensure that lamellocyte differentiation in the posterior lobes had not occurred earlier, we analyzed LG at day 2 post-parasitism. Again we found lamellocytes in the anterior but not posterior lobes (Figure S3C). This suggests that lamellocyte differentiation begins first in the anterior and that primary lobe cells are released into circulation. Furthermore, we analyzed *MSNF9-mCherry* larvae at day 3 post-parasitism and observed a similar pattern as in *Canton-S*, with frequent histolysis in anterior (13/17) but not in posterior (1/16 and 0/17) lobes (Figure 4C, D). Also, the induction of *MSNF9-mCherry* was much stronger in the anterior lobes while most *MSNF9-mcherry*^*+*^ cells present in the posterior lobes remained round and did not exhibit lamellocyte-like morphology, indicating a weaker response of PP (Figure 4C, D).

### Upregulation of STAT activity in posterior progenitors upon wasp parasitism

Our analysis indicates that PP resist differentiation following immune challenge by bacteria or wasp (Figure 4E). Downregulation of JAK-STAT signaling in the MZ following wasp parasitism is essential for the differentiation of blood cell progenitors into lamellocytes in the primary lobes (Makki et al., 2010). In contrast, wasp parasitism leads to an activation of JAK-STAT signaling in circulating hemocytes (and in somatic muscles) (Yang et al., 2015) and unrestrained activation of the JAK-STAT pathway in circulating hemocytes promotes lamellocyte differentiation (Bazzi et al., 2018). Yet, the role of JAK-STAT signaling in lamellocyte differentiation in the posterior lobes has not been assessed. Hence we investigated the status of JAK-STAT signaling in the whole LG to gain potential insight into the molecular basis for differential regulation of lamellocyte formation from anterior to posterior.

Accordingly, we used the 10xSTAT-GFP reporter (Bach et al., 2007) to analyze STAT92E activation in time-matched control (unparasitized) and parasitized larvae at day 2, 3 or 4 post-parasitism. In control larvae, 10x-STAT-GFP expression was essentially restricted to the posterior part of the anterior lobes at 120h (2 days post-parasitism) and 144h (3 days post-parasitism) AEL (Figure 5A, C). However it was barely detectable at later stage (4 days post-parasitism/ 168h AEL) (Figure 5E). In contrast, we observed consistent 10xSTAT-GFP expression throughout the posterior lobes from 120h to 168h AEL in parasitized larvae. Consistent with previous report (Makki et al., 2010), STAT92E activity was repressed in the anterior lobes following *L*. *boulardi* infection, as judged by the quasi absence of 10xSTAT-GFP expression in parasitized larvae (Figure 5B, D, F). Strikingly though, STAT92E activity was not repressed in the posterior lobes of parasitized larvae but was even highly upregulated at days 3 and 4 in comparison to unparasitized age-matched control lymph glands (Figure 5C-F).

**Figure 5:**
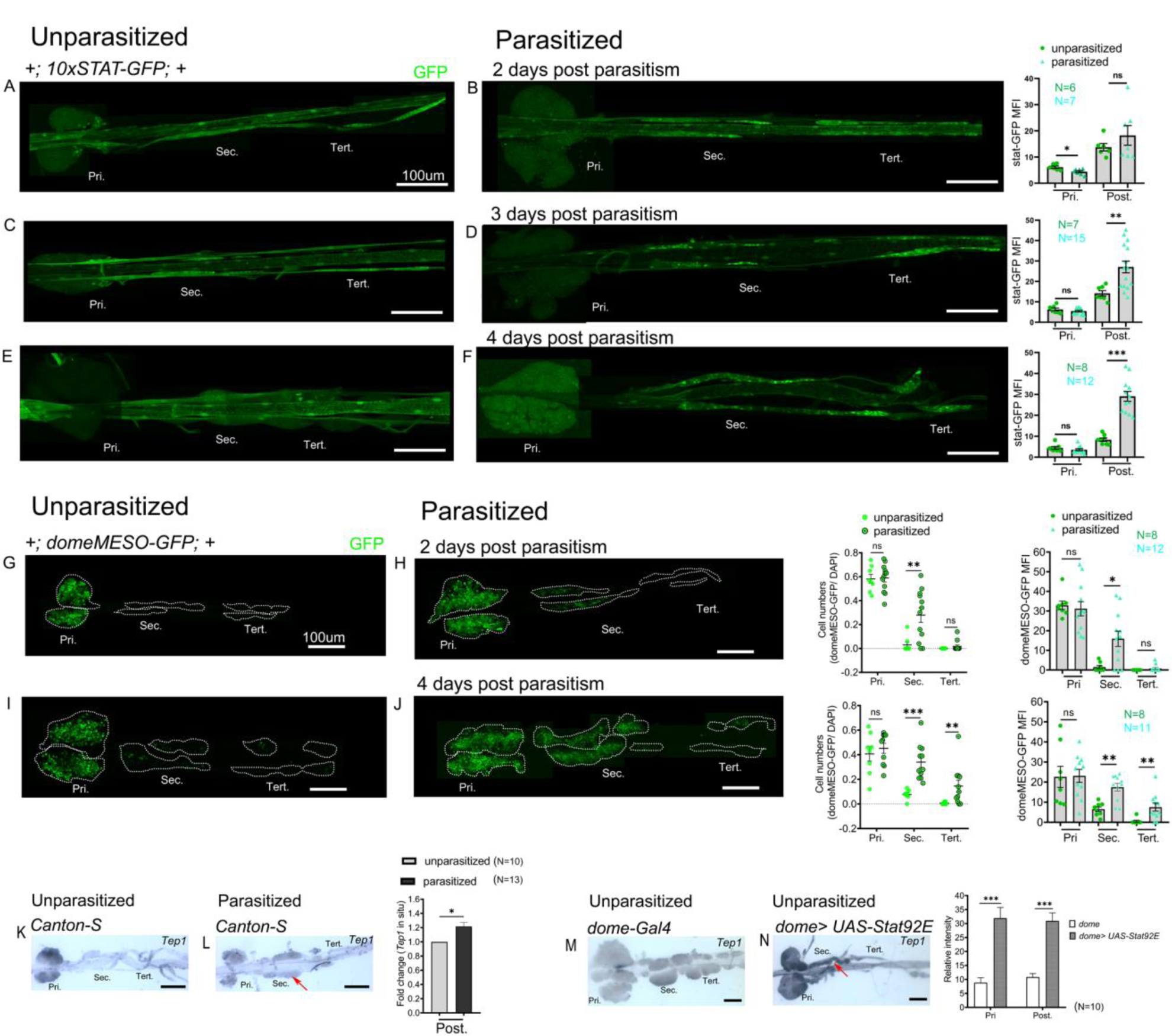
Posterior progenitors activate STAT in response to wasp parasitism. JAK-STAT activity is monitored using *10xSTAT-GFP* (A-F) or *domeMESO-GFP* (G-J) reporter. Reporter activity in the lymph gland lobes is compared and quantified between age-matched unparasitized and parasitized larvae at the indicated time points. (A, B) 2 days post-parasitism *10xSTAT-GFP* reporter activity (green) decreases in the primary lobes and remains unchanged in the posterior lobes. (C-F) 3 days post-parasitism and 4 days post-parasitism *10xSTAT-GFP* activity (green) increases in the posteriorlobes. Graphs representing the mean-fluorescence intensity (MFI) for *10xSTAT-GFP* are plotted for each time point. Mann-Whitney nonparametric test was used for statistical analysis and error bars represent SEM. (G, H) 2 days post-parasitism *domeMESO-GFP* reporter activity (green) remains unchanged in the primary lobes. (I, J) 4 days post-parasitism, domeMESO-GFP activity (green) significantly increases in the posterior lobes. Graphs representing ratio of *domeMESO-GFP*^+^/ DAPI^+^ cells and mean fluorescence intensity (MFI) are plotted for each time point. Mann-Whitney nonparametric test was used forstatistical analysis and error bars represent SEM. (K, L) RNA *in situ* hybridization shows that *Tep1* expression is induced in the posterior lobes (red arrow) post-parasitism as compared to unparasitized larval lymph glands. (M, N) Overexpression of *Stat92E* leads to transcriptional induction of *Tep1* (red arrow) in unparasitized conditions. Student’s t-test was used for statistical analysis and error bars represent SEM. *<*p*0.05, ** *p*<0.01, *** *p*<0.001 and ns indicates non-significant. Pri. indicates primary lobes, Sec. indicates secondary lobes, Tert. indicates tertiary lobes and Post. indicates posterior lobes. (A-N) scale bars: 100μm.

However, posterior lobes of parasitized 10xSTAT-GFP larvae appeared thinner than parasitized posterior lobes of wild type larvae. To rule out any anomaly due to this genetic background and confirm STAT92E activation, we analyzed domeMESO-GFP, another reporter of activated STAT (Hombria et al., 2005, Louradour et al., 2017). The 10xSTAT-GFP reporter consists of 5 tandem repeats of a 441-bp fragment from the intronic region of *Socs36E* with 2 STAT92E binding sites (Bach et al., 2007), whereas the domeMESO construct contains a 2.8-kb genomic fragment spanning part of the first exon and most of the first intron of *dome* (Hombria et al., 2005). Previous studies report the downregulation of *domeMESO*-lacZ 30h post-parasitism in the primary lobe (Makki et al., 2010). We analyzed *domeMESO*- GFP reporter 2 days and 4 days post-parasitism but surprisingly did not observe reduction in GFP signal in the primary lobes at any of these time points (Figure 5G-J). Persistent signal post-parasitism could be due to GFP perdurance. Nevertheless, secondary lobes showed significantly high GFP^+^ cells 2 days post-parasitism and at 4 days post-parasitism secondary as well as tertiary lobes had increased GFP- expressing cells (Figure 5G-J).

To gain further evidence for increased activity of STAT92E in the posterior lobes of parasitized larvae, we also assessed the expression of the STAT target *Tep1*. Indeed JAK overactivation as well as *L*. *boulardi* infection were shown to induce the expression of this opsonin (Lagueux et al., 2000, Wertheim et al., 2005, Salazar-Jaramillo et al., 2017). RNA *in situ* hybridization revealed a basal level expression of *Tep1* in the primary and posterior lobes in the wild type unparasitized larval lymph glands (Figure 5K). Upon parasitism, *Tep1* expression increased in the posterior lobes but not in primary lobes although most have disintegrated (Figure 5L). In addition, in line with the idea that *Tep1* is a target of the JAK/STAT pathway (Lagueux et al., 2000), we found that S*tat92E* overexpression in the progenitors induces *Tep1* expression in unparasitized conditions in the anterior as well as in the posterior lobes (Figure 5M, N).

The JAK-STAT pathway participates in various kinds of stress responses. Hence the activation of STAT92E in the posterior lobes may not be specific to wasp parasitism but could be a generic response to immune stress. To test this hypothesis, we infected larvae with *Pseudomonas entemophila*, a naturally occurring pathogen that induces a systemic immune response (Vodovar et al., 2005), or inserted a human hair in the hemocoel, which triggers lamellocyte differentiation (Lanot et al., 2001). However, these challenges did not cause activation of the STAT pathway in the lymph gland posterior lobes as assessed with the 10xSTAT-GFP reporter (Figure S4).

In sum, our results indicate that STAT92E activation in PP is likely a localized immune response to specific systemic cues triggered by wasp parasitism and suggest the existence of different mechanisms for regulating STAT92E in AP and PP.

### STAT92E limits the differentiation of lamellocytes in the posterior progenitors

Post-parasitism downregulation of JAK-STAT signaling is implicated in the differentiation of lamellocytes in the primary lobes (Makki et al., 2010). Our analysis thus far suggests that high levels of JAK-STAT signaling in the posterior lobes underlie their different response to parasitism. To test if high levels of JAK-STAT signaling inhibit lamellocyte induction in the posterior lobes, we knocked-down *Stat92E by RNAi* in the LG. The knockdown of *Stat92E* in the progenitors using *dome-Gal4* (or *tep4-GAL4*) at 25°C did not affect the survival of control larvae but caused lethality in parasitized larvae (Figure S5A, B), indicating STAT92E activation could be an essential part of the immune response. However, we obtained parasitized escapers when the larvae were raised at 21°C. In these conditions, STAT92E knock-down larvae exhibited stronger responses than controls in the anterior and posterior lobes, as judged by lamellocyte differentiation and fused/coalesced cells (Figure 6A-D). Notably, 60% (6/10) of STAT92E knockdown larvae show medium phenotypes against 20% (2/10) in controls. Additionally, the posterior lobes showed a decrease in progenitors as observed by reduced expression of *dome>GFP* (Figure6C-D), in agreement with our hypothesis that STAT92E blocks PP differentiation post-parasitism.

**Figure 6:**
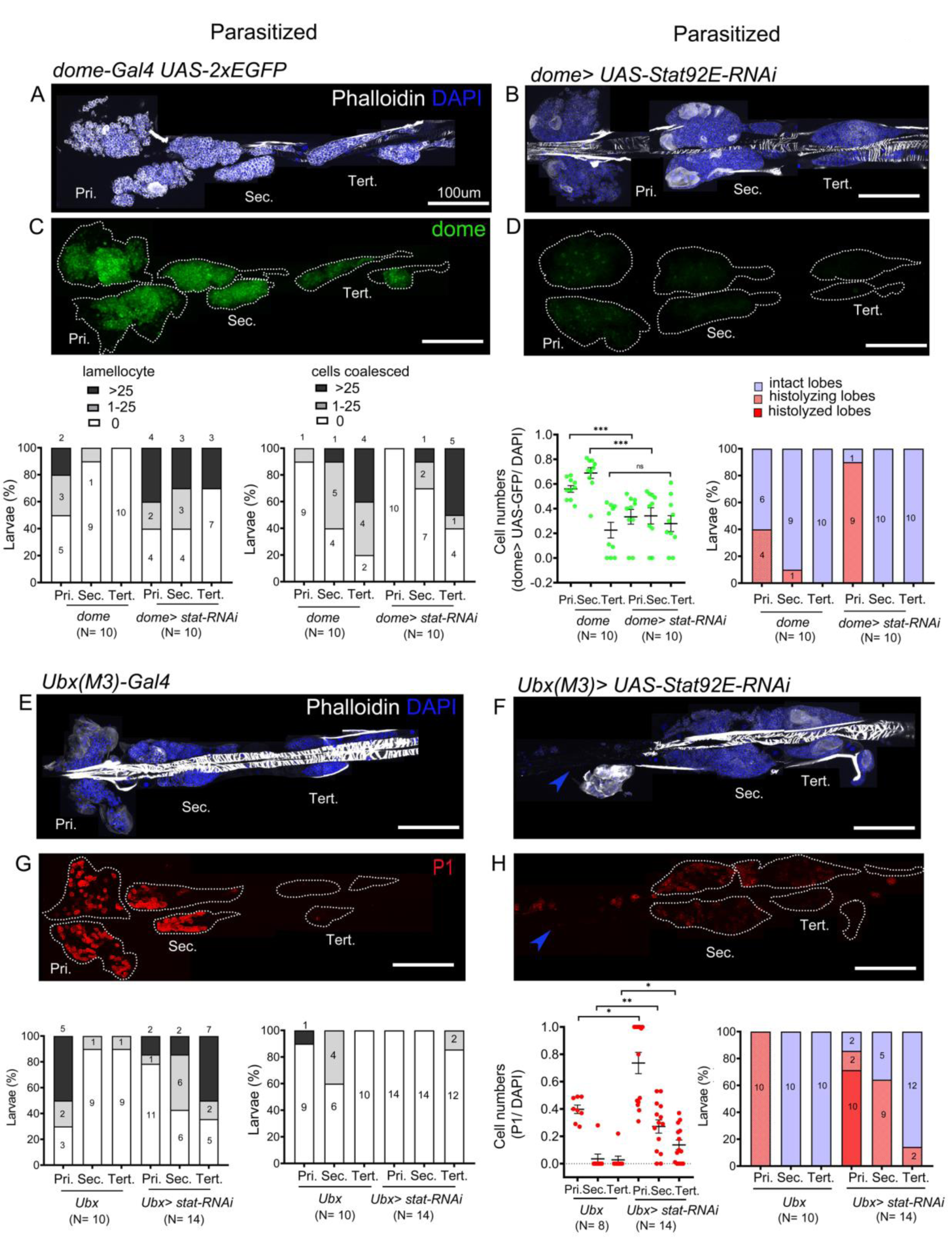
STAT limits lamellocyte differentiation in the posterior lobes. Lymph glands were analyzed 4 days post-parasitism, age-matched parasitized larvae were used for comparison. (A-H) Phenotypes for targeted knockdown of S*tat92E* in the progenitor population are shown. (A-D) S*tat92E* knockdown in the progenitors is associated with massive lamellocyte differentiation in the posterior lobes as revealed by phalloidin staining (white) (A, B) and a reduction in the progenitor pool as indicated by *dome-Gal4,UAS-2xEGFP* (green) expression (C, D) in parasitized larvae. (E-H) PP specific knockdown of S*tat92E* induces lamellocyte and plasmatocyte differentiation in the posterior lobes following parasitism. Graphs indicate quantification for the phenotypes-lamellocyte induction, compromised cell boundaries, dome^+^ or P1^+^ relative cell number, and lobe histolysis. (F, H) blue arrowheads indicate histolyzed lobes. Mann-Whitney nonparametric test was used for statistical analysis of dome and P1 and error bars represent SEM. \**p<0*.*05*, ** *p*<0.01, *** *p*<0.001 and ns indicates non-significant. Pri. indicates primary lobes, Sec. indicates secondary lobes, Tert. indicates tertiary lobes. (A-H) scale bar: 100µm.

As described above, the *Ubx(M3)-Gal4* is expressed in the PP but not in the primary lobes. Therefore we took advantage of this driver to specifically knockdown *Stat92E* in the posterior progenitors. Upon parasitism, more larvae showed lamellocyte induction in the posterior lobes in STAT92E knockdown conditions than in controls: 93% (13/14) of *Ubx(M3)-Gal4>STAT92E RNAi* larvae show medium phenotype versus 20% (2/10) in *Ubx(M3)-Gal4* controls. Conversely, only 7% (1/14) of *Stat92E* knockdown larvae displayed no phenotype, as compared to 50% (5/10) of control larvae. Additionally, we observed that many PP differentiate to plasmatocytes upon knockdown of *Stat92E* (Figure 6E-H). Of note, in unparasitized conditions, STAT92E knockdown using *dome-Gal4* or *Ubx(M3)-Gal4* does not trigger lamellocyte or plasmatocyte differentiation (Figure S5C, D, G-J) and *dome>GFP* expressing progenitors are maintained (Figure S5E, F).

These results strongly suggest that JAK-STAT signaling is essential for maintaining the progenitor pool and restricting lamellocyte differentiation post-parasitism. Our analyses reveal functional compartmentalization in the lymph gland progenitor pool which responds differentially to immune stress along the anterior-posterior axis. To unravel mechanisms regulating this compartmentalization we examined the status of key JAK-STAT pathway components in the whole LG.

### Differential regulation of the JAK-STAT inhibitor *latran* in lymph gland progenitors

Three ligands, Unpaired (Upd), Upd2, and Upd3 bind to the receptor Dome and trigger JAK-STAT signaling but only Upd3 is expressed in the lymph gland (Makki et al., 2010), both in the anterior and the posterior lobes (Table S1). Moreover *upd3* is required for JAK-STAT activation in the anterior lobes and its expression is downregulated 4h after wasp infestation (Makki et al., 2010). In agreement with a previous report (Jung et al., 2005), we found that the reporter line *upd3-GAL4* (Agaisse et al., 2003) is expressed in the MZ of the anterior lobes, as reported for *upd3* transcript (Makki et al., 2010), but also in the posterior lobes. Upon wasp parasitism its expression level was reduced in primary lobes but not in the posterior lobes (Figure 7A, B), even though they exhibit increased STAT92E activity at that time (see above). We also assessed the expression of the JAK-STAT pathway receptor *dome* using the *dome-GAL4* reporter (Bourbon et al., 2002, Jung et al., 2005). *dome-Gal4* is expressed in AP and PP in unparasitized larvae and, consistent with the previous report (Krzemien et al., 2007), we observed that it is down- regulated in the AP following parasitism (Figure 7C, D). Yet, no change was detectable in the PP. These results suggest that parasitism-induced activation of the JAK-STAT pathway in PP is not due to regulation of *upd3* or *dome* expression but other controls may operate.

**Figure 7:**
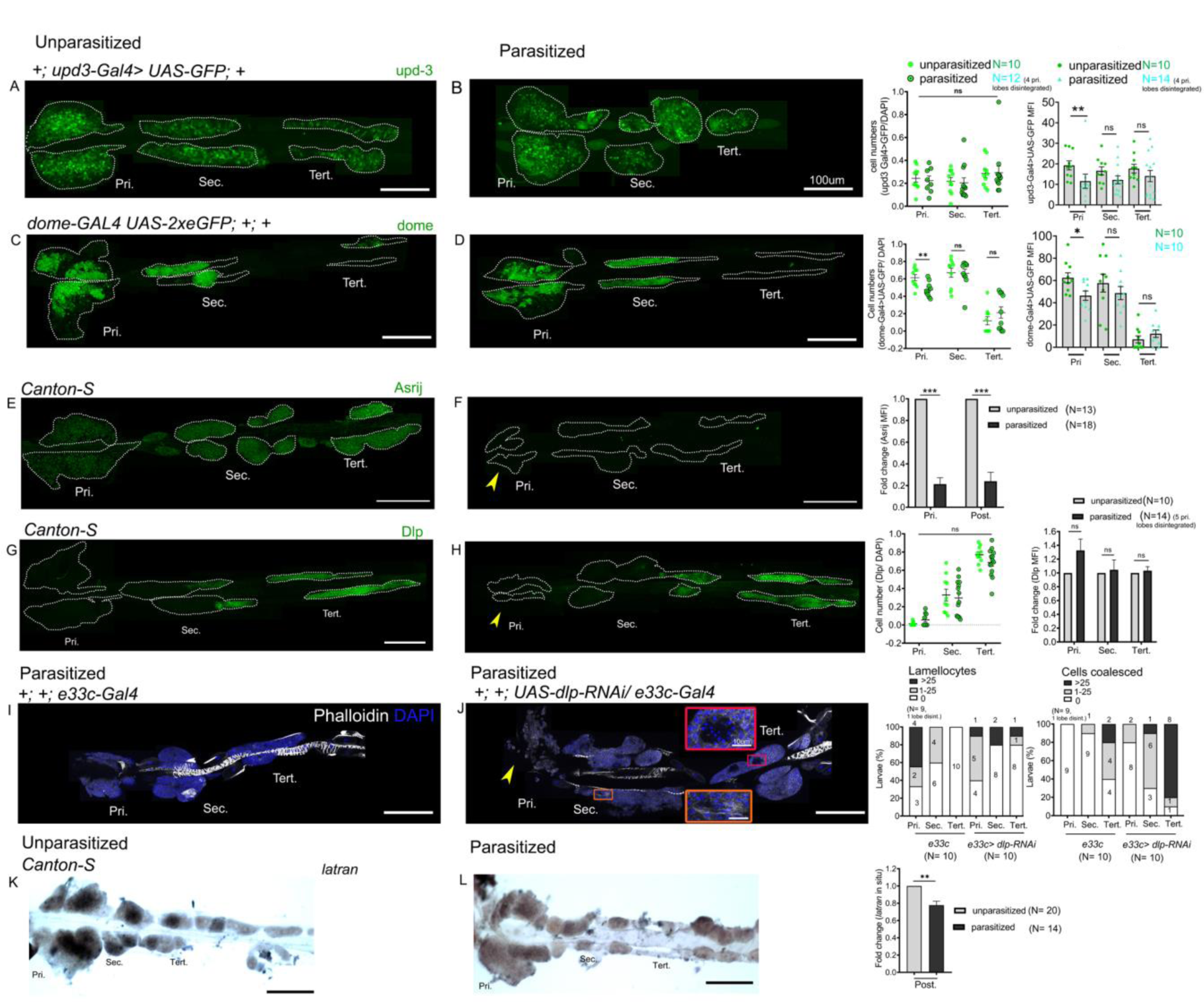
Differential regulation of the JAK-STAT inhibitor *latran* in AP and PP following parasitism. ymph glands were analyzed 4 days post-parasitism, age-matched unparasitized larvae were used for analysis in each case. (A, B) *upd3-Gal4> GFP* (green) expression reduces in the primary lobe and remains unchanged in the posterior lobes in parasitized conditions. *dome-Gal4>2xEGFP* (green) expression in unparasitized (C) and parasitized (D) conditions show reduced expression of *dome* in the primary lobes post-parasitism. Graphs represent ratio of cells positive for each marker, mean fluorescence intensity (MFI) in the respective lobes. Mann-Whitney nonparametric test was used for statistical analysis. (E, F) show reduced expression of Asrij (green) in the LG post-parasitism. Graphs represent fold-change for mean fluorescence intensity in the respective lobes; Student’s t-test is used for statistical analysis. Dlp (green) expression remains unchanged in unparasitized (G) and parasitized (H) conditions. Graphs represent ratio of Dlp positive cells and mean fluorescence intensity (MFI) in the respective lobes. Mann-Whitney nonparametric test and Student’s t-test was used for statistical analysis. (I, J) Targeted knockdown of *dlp* leads to lamellocyte formation and affects cell boundaries in the posterior lobes. Phalloidin (white) marks actin and is used for identifying lamellocytes and changes in cell morphology. (J) Orange inset shows lamellocyte formation in the secondary lobes and red inset displays compromised cell boundaries in the tertiary lobes. (K, L) RNA *in situ* hybridization for *latran* shows decreased expression in the PP post-parasitism. Graphs represent fold-change; Student’s t-test is used for statistical analysis. (F, H, J) Yellow arrowhead indicates disintegrating primary lobes. Error bars represent SEM. \**p<0*.*05*, ** *p*<0.01, *** *p*<0.001 and ns indicates non-significant. Pri. indicates primary lobes, Sec. indicates secondary lobes, Tert. indicates tertiary lobes and Post. indicates posterior lobes. (A-L) scale bar: 100µm or 10µm in inset.

Along that line, we previously showed that the endosomal protein Asrij, a conserved modulator of stem and progenitor maintenance, positively regulates STAT92E activation in the lymph gland (Sinha et al., 2013) (Figure S6A, B). Hence we investigated whether increased STAT92E activation in the PP upon wasp parasitism could be a result of increased Asrij expression. However Asrij protein levels were significantly reduced in both anterior and posterior lobes upon wasp parasitism (Figure 7E, F) suggesting an *asrij*- independent mechanism of STAT92E activation post parasitism. In agreement with this, *asrij* null LGs still showed increased 10XSTAT-GFP reporter expression in the posterior lobes following parasitism (Figure S6C, D). In addition, we found that Dlp, an ECM component that promotes JAK-STAT signaling by stabilizing Upd (Hayashi et al., 2012, Zhang et al., 2013), is highly expressed in the posterior lobes (see above). It could thus participate in the differential regulation of the JAK-STAT pathway between AP and PP and/or in the regulation of PP fate. Immunostaining showed no change in Dlp levels post-parasitism and it remained high in posterior lobes (Figure 7G, H). Nonetheless, we tested the role of Dlp in the response to parasitism by knocking-down its expression by RNAi using the *e33c-Gal4* that shows widespread expression in the lymph gland lobes (Harrison et al., 1995, Kulkarni et al., 2011). Interestingly, the knock- down of *dlp* increased the occurrence of mild phenotypes in the posterior lobes as compared to control larvae, whereas it did not enhance the response to parasitism in the anterior lobes (Figure 7I, J). This suggests that Dlp, by promoting sustained JAK-STAT signaling in the posterior lobes, contributes to the different responses of the anterior and posterior lobes progenitors.

Finally, it was shown that downregulation of JAK-STAT signaling by Latran is required for lamellocyte differentiation in the primary lobes (Makki et al., 2010). RNA *in situ* analysis of unparasitized LG showed high levels of *latran* in the primary lobes as reported (Makki et al., 2010). Additionally we observe high levels of *latran* in the secondary lobes compared to tertiary lobes (Figure7K). Upon parasitism *latran* expression is reduced in the posterior lobes (Figure 7L), indicating that differential pathway activation may be regulated by Latran.

## Discussion

The *Drosophila* lymph gland has been used as a powerful system to study blood cell progenitor maintenance. However intra-population heterogeneity has not been explored previously. Here we introduce the lymph gland as a model to analyze progenitor heterogeneity at the level of the complete hematopoietic organ by phenotypic marker expression, proliferative capacity and differentiation potential. Firstly we show that posterior lobes consist of a significant population of progenitors and are essentially devoid of differentiated cells. PP express progenitor markers like *Tep4, dome, DE-cad* in the secondary lobes and in some cells of the tertiary lobe but very few cells expressing differentiation markers such as Pxn, P1/NimC1 or ProPO are present. Furthermore, while most cells of the posterior lobes express low levels of *col* like the anterior progenitors, part of the tertiary lobes express high levels of *col*, like the PSC in the anterior lobes. However we did not observe expression of other PSC markers such as *Antp* or *hh*, suggesting that there is no clear homologue of the PSC/niche in these lobes. While several autonomous factors and local signals emanating from the PSC have been implicated in progenitor maintenance in the anterior lobes (Banerjee et al., 2019), how progenitor fate is maintained in the posterior lobes remains largely unknown and certainly deserves further investigation. The identification of genes overexpressed in the PP, such as the transcription factors Apt or the ligand Net-B, paves the way for such investigations.

AP are responsive to systemic cues/ long range signals present in the hemolymph. For instance, they are sensitive to nutrient deprivation and olfactory cues, which regulate specific signaling pathways in the anterior lobes (Benmimoun et al., 2012, Shim et al., 2012, Tokusumi et al., 2012, Shim et al., 2013). It would be all the more interesting to study whether the same processes operate in PP as we revealed clear differences between AP and PP response to systemic cues elicited by immune challenges. Our finding that *5-HT1B* and *5-HT1A*, which code for serotonin receptors, are expressed at higher levels in the posterior lobes suggests that progenitor fate or function could be controlled by serotonin circulating through the hemolymph or produced by neighboring cells. Indeed, besides its function as a neurotransmitter, serotonin acts as a peripheral hormone and has an immunomodulatory effect on blood cells (Herr et al., 2017). Notably, mice deficient for the serotonin receptor 5-HT2B show altered bone marrow composition, with increased granulocyte precursors and reduced immature endothelial precursors (Launay et al., 2012). Moreover, phagocytosis is severely impaired in *Drosophila* mutants for serotonin receptors 5-HT1B and 5-HT2B (Qi et al., 2016).

Although our transcriptome analysis revealed a strong over-representation of genes associated with ECM organization (such as the laminins LanB1/B2/A, Collagen 4A1, Viking, Tiggrin, Glutactin or Papilin) among those overexpressed in the primary lobes, the HSPG Dlp and the ECM receptor Dg were overexpressed in the PP. Dlp was shown to regulate PSC size by regulating the response to the Decapentaplegic (Dpp) signaling pathway (Pennetier et al., 2012), but its function in the prohemocytes is unknown. Dg binds ECM proteins like Perlecan, whose mutation causes reduced lymph gland growth and premature differentiation of the progenitors in the primary lobes (Grigorian et al., 2013). Similarly, the ECM protein Tiggrin was found to control the progression of intermediate progenitors to mature plasmatocytes (Zhang and Cadigan, 2017). Grigorian et al. (2011) also suggested that hemocytes digest only a small part of the ECM to facilitate their dispersal and most of the ECM is left intact during metamorphosis. Systemic as well as local signals regulating ECM secretion and adhesiveness play an important role in vertebrate bone marrow to regulate blood cell quiescence, maintenance or egress. (Klamer and Voermans, 2014, Gattazzo et al., 2014, Zhang et al., 2019, Khadilkar et al., 2020). However in-depth analysis of the widely dispersed vertebrate hematopoietic compartment is technically challenging. A deeper understanding of the differential expression of ECM components in the *Drosophila* LG and how they regulate signaling in and maintenance of progenitors could reveal evolutionary conserved mechanisms that help understand bone *marrow* hematopoiesis.

Our results bring new insights into the origin and fate of the posterior lobes during development. Posterior lobes flank the dorsal vessel in the abdominal segments (Jung et al., 2005, Banerjee et al., 2019), however there is no literature regarding the ontogeny of the posterior lobes. In the embryo the homeotic gene *Ubx* provides positional cues to restrict the formation of the primary lobes to the thoracic segments (Mandal et al., 2004). Yet, our transcriptome show that *Ubx* is strongly expressed in the PP and we observe that Ubx is specifically expressed in the tertiary lobes of third instar larval lymph glands. Moreover, lineage tracing analysis with *Ubx(M3)-Gal4* strongly suggests that all the cells of the posterior lobes (and none of the anterior lobes) are derived from the Ubx^+^ anlage, *i*.*e*. the embryonic abdominal segments A1 to A5 (Lo et al., 2002). It would be interesting to study the role of Ubx and the more posterior Hox Abdominal-A (Abd-A) or Abdominal-B (Abd-B) (which are not expressed in the AP or PP) in providing positional cues for seeding the posterior lobes. Likewise several Hox, including Ubx ortholog HoxA7, have been implicated in HSPC development in mammals (Collins and Thompson, 2018). Whether they contribute to the emergence of distinct pools of blood cell progenitors during development is still unknown.

Lymph gland lobes histolyze at the onset of metamorphosis (Lanot et al., 2001, Grigorian et al., 2011, Makhijani et al., 2011, Gold and Bruckner, 2015). A previous study focused on the anterior lobes showed that secondary lobes express Pxn at 4hr APF but disperse before terminal differentiation (Grigorian et al., 2011). We observe that in all the posterior lobes, cells that persist till 10 or 15 hr APF still express progenitor markers and do not undergo terminal differentiation. Pupal and adult blood cells derive from the embryonic and lymph gland lineages (Holz et al., 2003). We speculate that under normal conditions PP act as a reserve/ long term pool of progenitors that could have specific functions in larval and/or pupal and adult stages. Along that line, it was proposed that undifferentiated blood cells derived from *col*-expressing posterior lobe hemocytes persist in the adult (Ghosh et al., 2015), but a recent study challenged these conclusions and found no evidence for active hematopoiesis during adulthood (Sanchez Bosch et al., 2019). However, a detailed lineage tracing analysis for understanding the contribution of AP and PP to pupal and adult stages has not been possible. It is anticipated that the present identification of new progenitor markers and of a GAL4 driver specifically expressed in the posterior lobes should help characterize the fate of lymph gland progenitors in the pupal and adult stages.

Importantly, we show here that the PP exhibit a distinct behavior as compared to AP and our data support the hypothesis that differential regulation of the JAK-STAT pathway underlies this functional compartmentalization. In response to wasp parasitism downregulation of JAK-STAT signaling is required for the differentiation of lamellocytes in the primary lobes (Makki et al., 2010, Louradour et al., 2017). Our analysis of the lymph gland in its entirety indicates that PP exhibit higher level of JAK-STAT signaling than the anterior lobes and further activate the pathway in response to wasp parasitism. Moreover, the maintenance of high level of STAT92E activity in the PP appears to be required to prevent lamellocyte differentiation in these cells. Interestingly, wasp infestation was also shown to cause JAK-STAT signaling activation in the larval somatic muscle, which is essential for mounting an efficient immune response (Yang et al., 2015). Thus wasp infestation seems to lead to differential regulation of the JAK-STAT pathway in multiple target organs. Notably, sustained JAK-STAT signaling in the PP, by contributing to the expression of complement like factor Tep1, could further activate the humoral arm of immunity. This also raises the possibility that under normal physiological conditions PP act as a reserve/ long term progenitor pool of hemocytes that are activated to play critical functions under specific immune conditions.

It is interesting to note that while infestation by the specialist wasp *L*. *boulardi* induces melanization and lamellocyte differentiation, venom from the generalist *L*. *heterotoma* is more immune suppressive and causes hemocyte lysis (Schlenke et al., 2007). Toll and JAK/STAT pathway genes are some of the most highly expressed genes in the anti-parasite immune response to *L*. *boulardi* leading to Tep1 upregulation, which is not seen in *L*. *heterotoma* infested flies. However upstream signals that bring about this regulation are not known.

Although the decrease in *latran* expression that we observe in the PP following wasp parasitism could strengthen JAK-STAT signaling, how this pathway is activated in the PP remains unclear. Integration of external stimuli to produce an appropriate response is essential for maintaining blood cell homeostasis and survival. Post-parasitism the involvement of cytokines Upd3 and Upd2 in activating JAK-STAT signaling in the larval muscles indicates the role of long-range signals in integrating response to immune stress (Yang et al., 2015). Post-parasitism we observe no change in the expression of Upd3 in the lymph gland. However, Dlp is known to promote JAK-STAT activation by binding Upd cytokines (Zhang et al., 2013) and it is expressed at high levels in the PP. Dlp could thus sequester Upd2 and/or Upd3 from the hemolymph, leading to the activation of JAK-STAT signaling in the PP. Enhanced expression of the ECM components, like the HSPG Dlp, in the posterior may help integrate cytokine signals leading to local activation of pathways such as JAK-STAT. Further the role of signals such as serotonin or developmental events remains to be investigated.

In summary, we show that the *Drosophila* larval hematopoietic organ, the lymph gland, has a heterogeneous pool of progenitors whose maintenance is spatially and temporally regulated and we reveal a previously unexpected role for JAK-STAT signaling in maintaining the posterior progenitors in the presence of immune challenge (Figure 8). The *Drosophila* lymph gland could serve as a potent model to understand blood cell progenitor heterogeneity and our findings pave the way for future investigation aimed at understanding the spatio-temporal regulation of progenitor fate in normal and pathological situations *in vivo*. Wasp infestation provides refined and relevant interventions, akin to a sensitized genetic background, to highlight the context-dependent roles of signaling and progenitors. Using infestation as a tool we could uncover differences in the progenitor response to systemic signals and evolutionarily conserved signaling pathways that regulate these. Understanding such interactions between the immune system and its stimuli could inform developmental analysis and our work sets the stage for future studies to decipher these.

**Figure 8:**
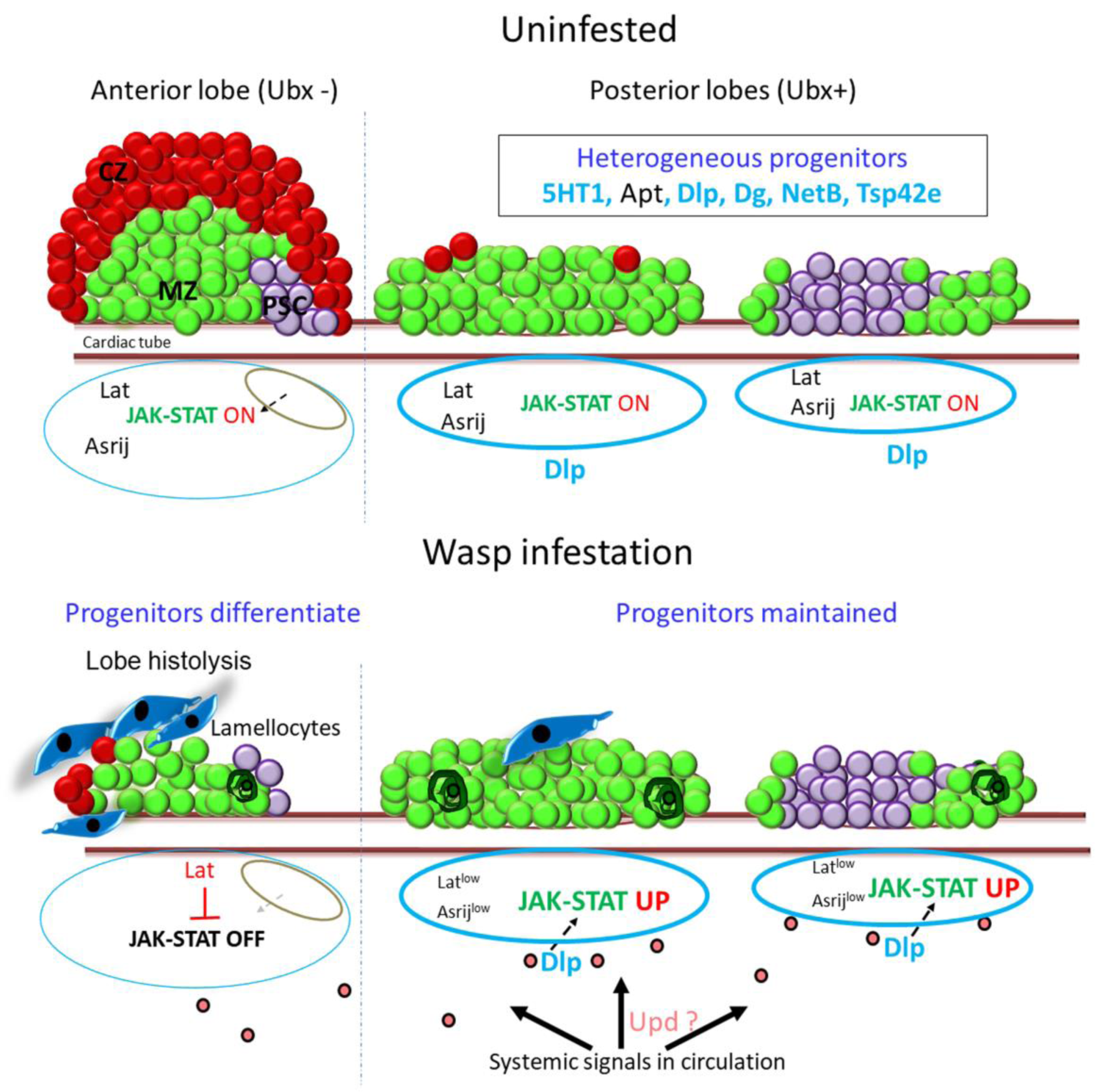
Model depicting lymph gland progenitor heterogeneity and response to immune challenge. Schematics showing the entire *Drosophila* third instar larval lymph gland. Uninfested - primary lobe is demarcated into the niche/ PSC (purple), progenitor/ MZ (green) and differentiated/ CZ (red) zones. Posterior lobes harbor a heterogeneous progenitor pool that is not segregated into zones. Posterior progenitor markers identified from this study are indicated in light blue (Extracellular ligands, matrix components and receptors) or black (transcription factors) font. The status of JAK-STAT signaling is indicated below for each lobe. Upon wasp infestation, JAK-STAT pathway is downregulated in anterior progenitors leading to lamellocyte differentiation (dark blue) and histolysis. Systemic signals lead to production of ligands such as Upd (pink) that may be selectively trapped by extracellular matrix components such as Dlp around the posterior lobes (light blue border), leading to increased JAK-STAT signaling, in an Asrij-independent and Latran-dependent manner. Posterior progenitors are maintained and few coalesced cells are seen (dark green). PSC: posterior signaling centre; MZ: medullary zone; CZ: cortical zone

## Acknowledgements

We thank the *Drosophila* Hemocyte Biology and European Drosophila Research Conference (EDRC) meeting participants for valuable discussions and feedback; *Drosophila* community for fly stocks and antibodies; National Centre for Biological Sciences, Fly Facility for stocks; JNCASR Imaging facility and our laboratory members for valuable inputs and suggestions. This work was funded by the Indo-French Centre for the Promotion of Advanced Research (IFCPAR/ CEFIPRA) grant to MI and LW; MI’s work was also supported by SERB grant, J C Bose award project and Jawaharlal Nehru Centre for Advanced Scientific Research. LW was supported by grants from the Agence Nationale pour la Recherche and Fondation ARC.

## Inventory of supplementary information

### Supplementary Figures

**Supplementary figure S1:**
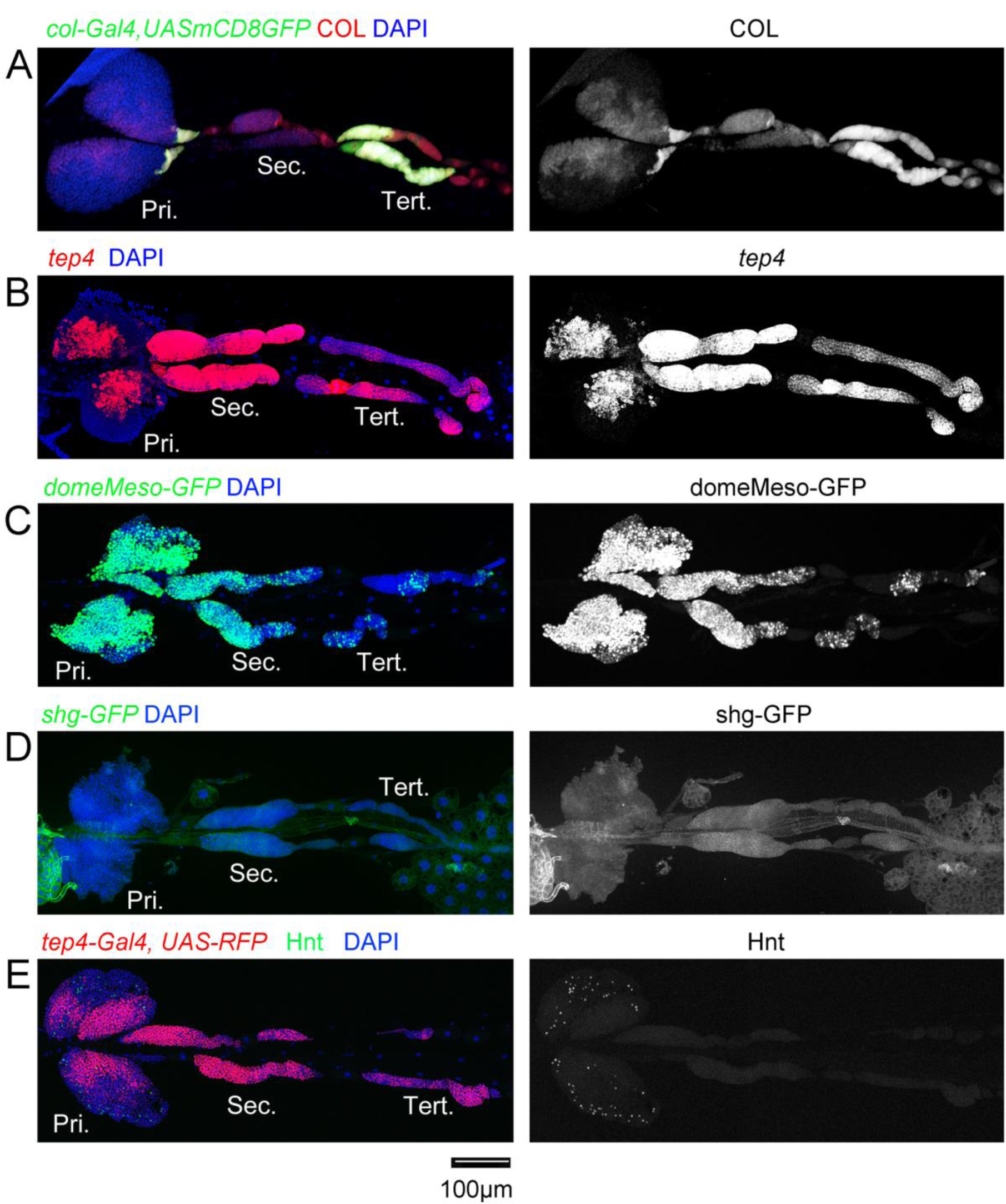
Lymph gland gene expression analysis. (A-E) Third instar larvae whole lymph gland preparations showing the expression of (A): *col-Gal4,UAS-GFP* (green) and Col (red), (B) *Tep4* (red), (C) *domeMESO-GFP* (green), (D) *shg-GFP* or (E) *Tep4-Gal4,UAS-RFP* (red) and Hnt (green) as revealed by fluorescent immunostaining (A, C-E) or *in situ* hybridization (B). Nuclei were stained with DAPI (blue). Pri. indicates primary lobes, Sec. indicates secondary lobes, Tert. indicates tertiary lobes. Right panels: single channel images. Scale bar: 100µm.

**Supplementary figure S2:**
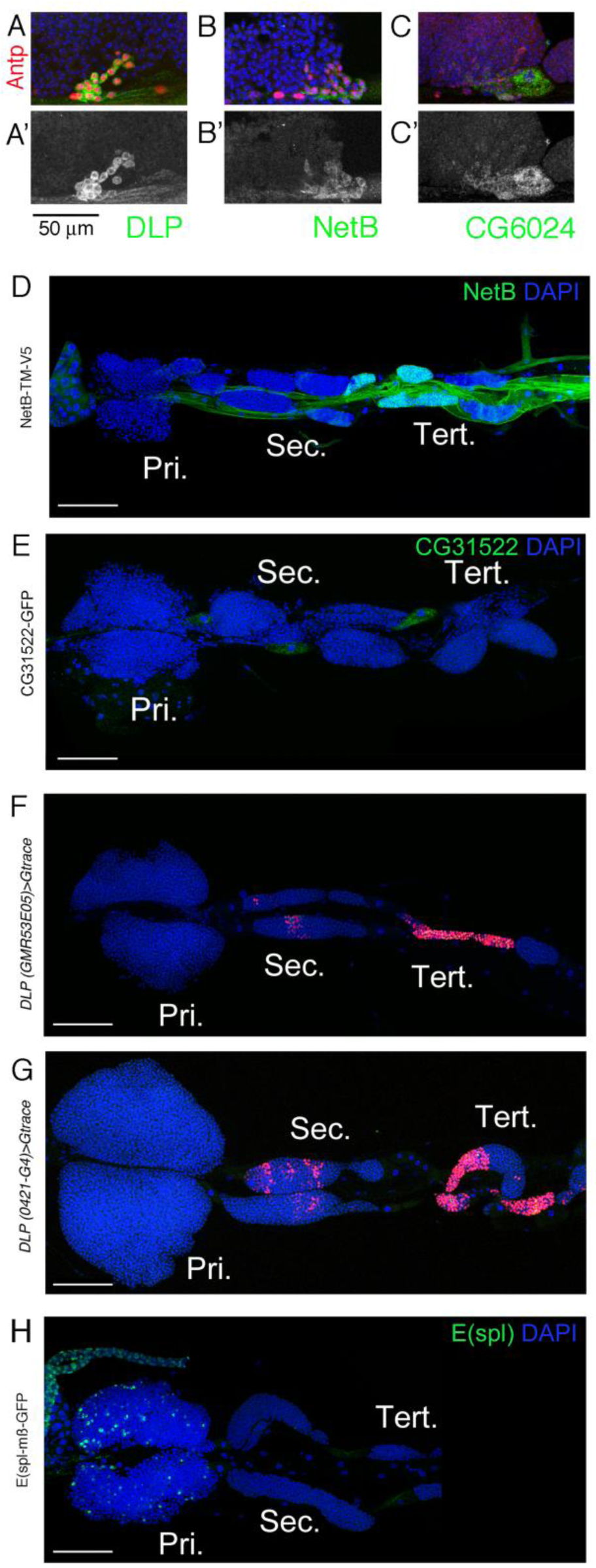
Expression of new markers in the lymph gland. (A-C) High magnification views of the posterior tip of the anterior lobe in third instar larval lymph glands show that Dlp (A, green), NetB-GFP (B, green) and CG6024-GFP (C, green) are expressed in the PSC, as revealed by Antp immunostaining (red). (A’-C’) show the green channel only. Scale bar: 50μm. (D-H) Third instar larvae whole lymph gland preparations showing the expression of NetB-TM-V5, CG31522-GFP, E(spl)mß-HLH-GFP or dlp enhancers GMR35E05 and 0421-G4 as revealed by immunostaining against V5 (D, green) or GFP (E,H, green) or by G-trace (F,G, red). Pri. indicates primary lobes, Sec. indicates secondary lobes, Tert. indicates tertiary lobes. (D-H) Scale bar: 100μm. (A-H) Nuclei were stained with DAPI (blue).

**Supplementary figure S3:**
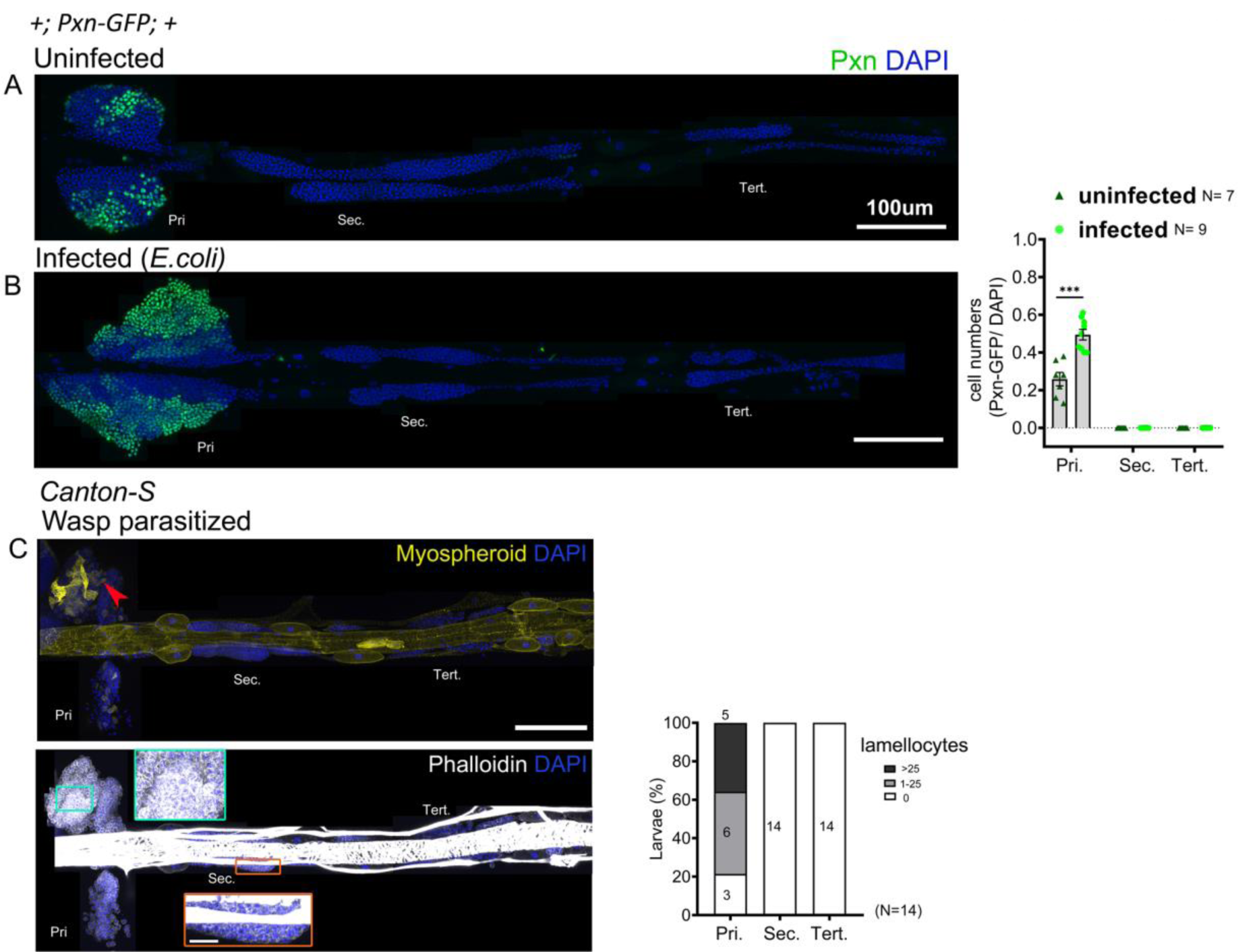
PP do not differentiate in response to *E*. *coli* infection and wasp parasitism. (A, B) Primary lobes show increased expression of *Pxn-GFP* (green) in response to *E*. *coli* infection whereas PP are maintained. Graphs indicate ratio of Pxn^+^/ DAPI^+^ cells. Mann-Whitney nonparametric test was used for statistical analysis and error bars represent SEM. (C) Lymph gland response 2 days post-parasitism by *L*. *boulardi*. Immunostaining against the ß-integrin Myospheroid (yellow) reveal lamellocyte differentiation (red arrowhead). Phalloidin (white) marks actin. Green and orange inset: high magnification views of the primary and secondary lobes, respectively. Pri. indicates primary lobes, Sec. indicates secondary lobes and Tert. indicates tertiary lobes. Graphs indicate quantification for lamellocyte induction. \**p<0*.*05*, ** *p*<0.01, *** *p*<0.001 and ns indicates non-significant. Scale bar: 100µm.

**Supplementary figure S4:**
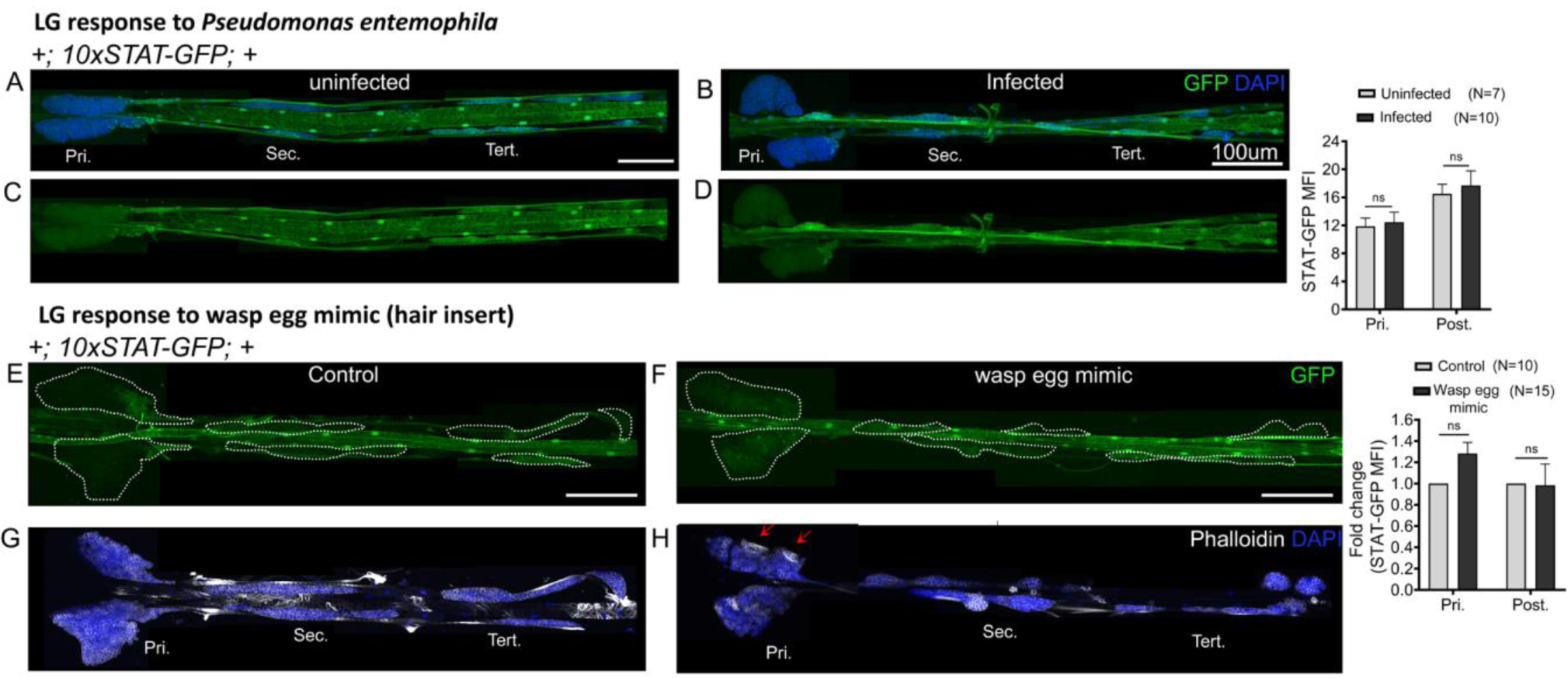
*10xSTAT-GFP* reporter activity in response to bacterial infection and wasp egg mimic. (A-D) *10xSTAT-GFP* reporter activity (green) remains unchanged upon *Pseudomonas entemophila* infection. Uninfected larval lymph gland (A, C) compared with *Pseudomonas entemophila* infected lymph gland (B, D). Mann-Whitney nonparametric test was used for statistical analysis. (E-H) *10xSTAT-GFP* reporter (green, E, F) and Phalloidin (white, G, H) in control (E, G) and wasp egg mimic conditions (F, H) show no change in reporter activity. Lamellocytes in the primary lobes are shown (red arrows). Graphs represent mean-fluorescence intensity (MFI) for *10xSTAT-GFP*. Student’s t-test is used for calculating fold change, error bars represent SEM. \**p<0*.*05*, ** *p*<0.01, *** *p*<0.001 and ns indicates non-significant. Pri. indicates primary lobes, Sec. indicates secondary lobes, Tert. indicates tertiary lobes and Post. indicates posterior lobes. Scale bar: 100µm.

**Supplementary figure S5:**
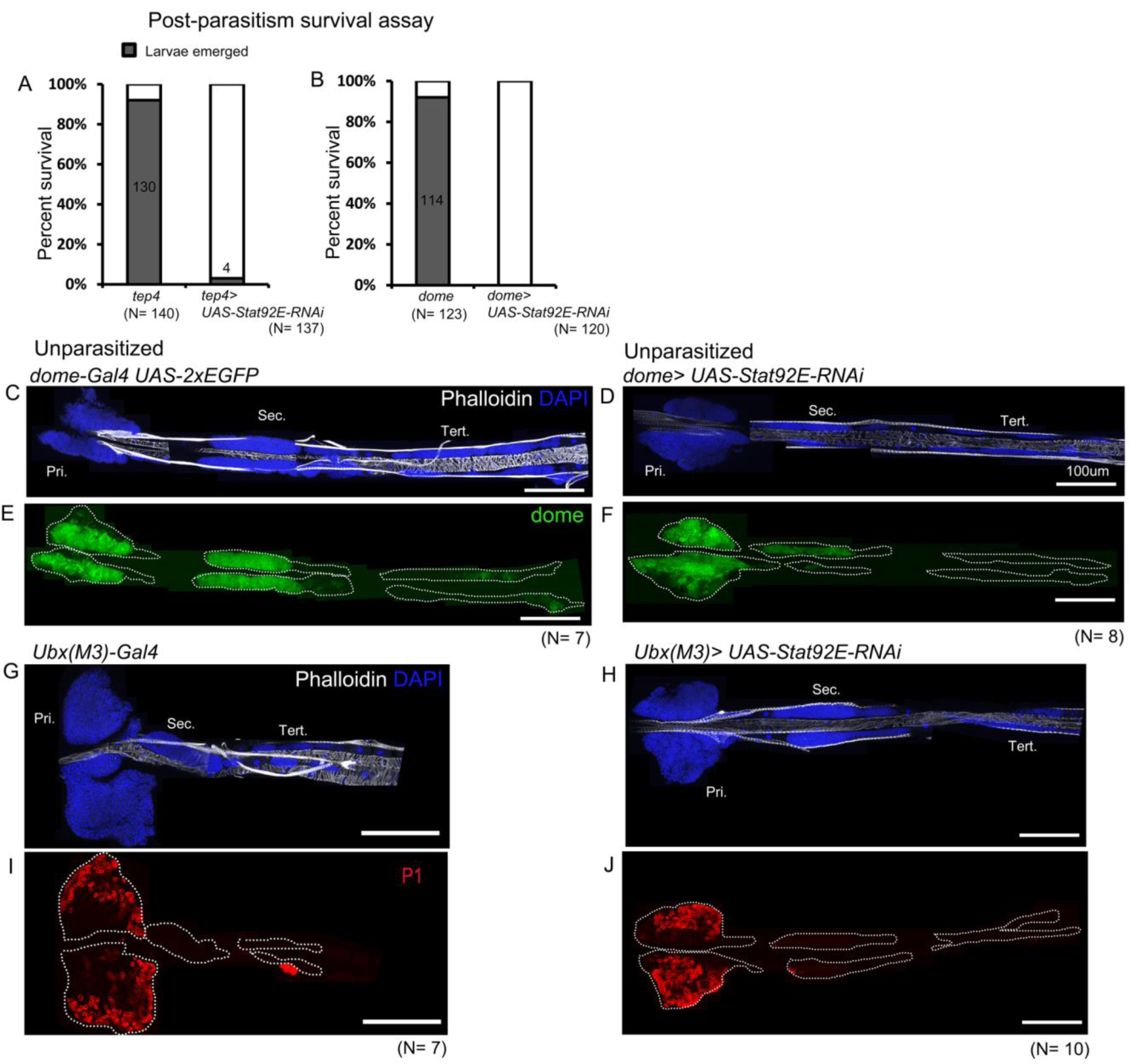
*Stat92E* is essential for survival post-parasitism. (A, B) Percentage of survival in S*tat92E* perturbed conditions post-parasitism. (C-J) Targeted knockdown of S*tat92E* in unparasitized conditions is shown. (C-F) S*tat92E* knockdown in the progenitors does not lead to lamellocyte differentiation as revealed by phalloidin staining (white) (C, D) and does not affect the progenitor pool as indicated by *dome-Gal4,UAS-2xEGFP* (green) expression (E, F). (G-J) PP specific knockdown of S*tat92E* does not induce lamellocyte and plasmatocyte differentiation in the posterior lobes. Pri. indicates primary lobes, Sec. indicates secondary lobes and Tert. indicates tertiary lobes. Scale bar: 100µm.

**Supplementary figure S6:**
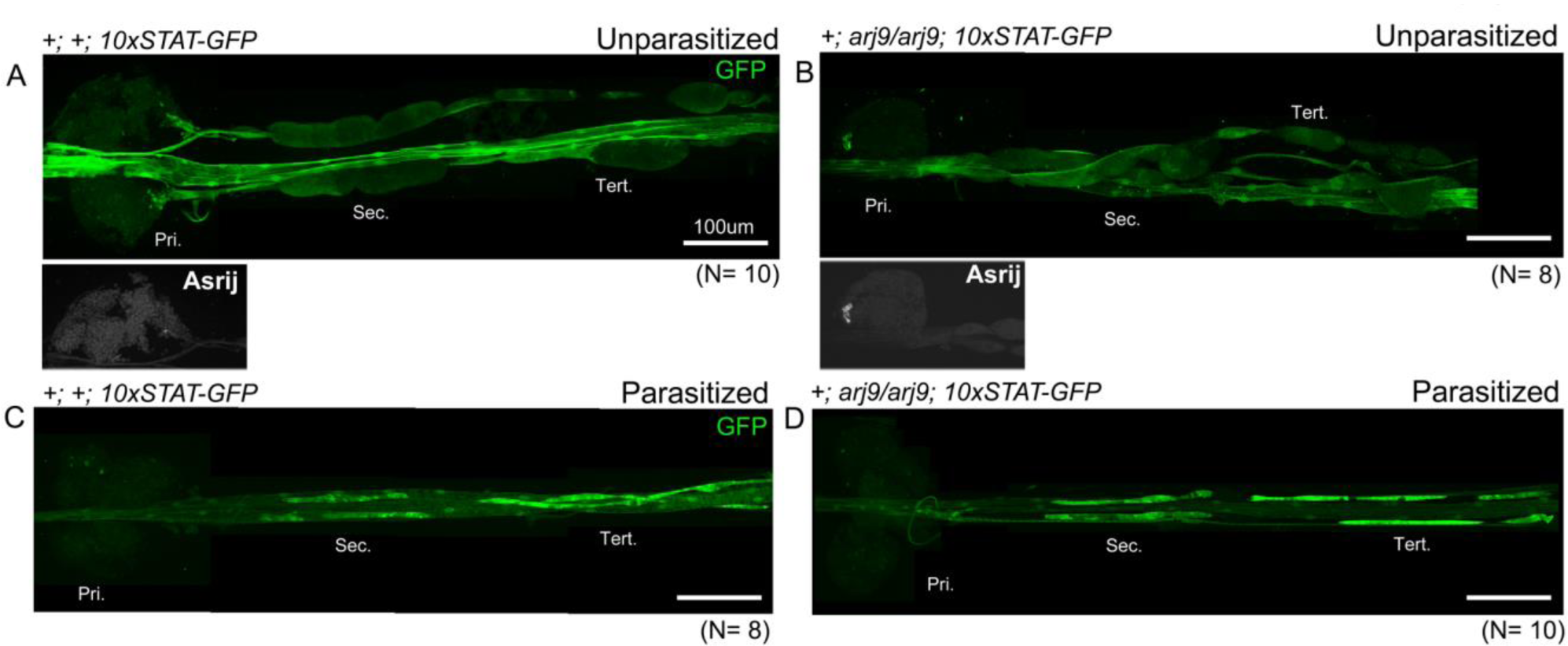
Asrij-independent upregulation of STAT activity in posterior progenitors following parasitism. (A, B) *asrij* null mutants show reduced expression of *10xSTAT-GFP* in the posterior lobes. (C, D) *10xSTAT-GFP* activity increases in *asrij* null mutants following *L*. *boulardi* parasitism as compared to unparasitized controls. Pri. indicates primary lobes, Sec. indicates secondary lobes and Tert. indicates tertiary lobes. Scale bar: 100µm.

### Supplementary Tables

**Supplementary Table S1. List of genes expressed in the lymph gland anterior and/or posterior lobes**

The expression level of each gene (RPKM values) in each of the 6 RNA-seq samples (3 anterior lobes, 3 posterior lobes) is indicated. Only genes with a FPKM≥1 in all 3 samples of the anterior lobes or of the posterior lobes are considered.

**Supplementary Table S2. List of differentially expressed genes between posterior and anterior lobes (p-value<0.01, fold change >1.5)**

Blue: genes over-expressed in the posterior lobes. Red: genes over-expressed in anterior lobes.

**Supplementary Table S3. Over-represented GO categories (Biological Processes, Cellular Components, Molecular functions) among genes over-expressed in the anterior or posterior lobes**

